# 3’ to 5’ Translation of Circular RNAs?

**DOI:** 10.64898/2025.12.08.692888

**Authors:** Zhe Lin, Xiangyou Pi, Ziwei Lv, Tianliang Liu, Huajun Yin, Danni Shi, Muxi Li, Juan Du, Yanchao Yang, Shiyu Wang, Peng Wang, Yangmei Qin, Wenbo Lin, Tao Tao, Ling Sun, Wenfeng Chen, Xi Zhang, Yufeng Yang, Zhiliang Ji

## Abstract

For decades, the 5’→3’ direction of translation has been a central dogma of molecular biology(*1*). Here we present evidence that eukaryotic circular RNAs (circRNAs) can serve as templates for 3’→5’ backward translation (BT), yielding polypeptides with distinct sequence and structural features not found in canonical proteomes. Mining of more than 6,000 multi-omics datasets identified ∼1 million candidate BT proteins across eukaryotes, including 59,000 high-confidence human BT proteins supported by mass spectrometry. Genetic combinatorial experiments (KO and KI) and cell-free translation of synthetic circRNAs establish BT as a conserved mechanism. Loss-of-function studies of *hs.*circCAPN15 in human and *dm.*circROLS in *Drosophila* underscore the functional importance of BT-derived proteins. Our work challenges the long-standing unidirectional translation paradigm, expands the functional landscape of the genetic code, and reveals a hitherto hidden layer of proteomic complexity with broad implications for biology and therapeutics.

## Main Text

Delimiting the molecular diversity space serves as a fundamental prerequisite for probing biological complexity. While the telomere-to-telomere (T2T) human genome assembly (v2.0) now provides the most complete genomic reference(*2*), annotating 136,194 coding sequences (CDSs) (GRCh38.p14), proteome discovery has stagnated. The proteome discovery rates have declined sharply, with novel human protein identifications in UniProt dropping from nearly 2,000 (2010–2015) to less than 500 (2016-2023)(*3*). Paradoxically, a substantial proportion of mass spectra in human MS data remains unmatched to proteins in reference databases(*4*), such as UniProtKB(*5*) and NCBI Refseq(*6*), despite deploying artificial intelligence-based methods to advance MS data deciphering(*7*). This paradoxical situation indicates either (i) limitations in current annotation paradigms, or (ii) the existence of non-canonical translation products undetected by standard pipelines.

Emerging evidence suggests that non-coding RNAs (ncRNAs), including long ncRNAs (lncRNAs) and circular RNAs (circRNAs), can be translated into functional small proteins or polypeptides(*8–10*). These ncRNA-derived peptides contribute to a subset of unassigned spectra in mass spectrometry data(*11*). Unlike linear RNAs, circRNAs form covalently closed loops through back-splicing and lack a 5’ 7-methylguanosine (m^7^G) cap, challenging conventional models of protein synthesis(*12*). Current findings indicate that circRNA translation proceeds via cap-independent mechanisms, potentially facilitated by internal ribosomal entry site (IRES)-like elements, N6-methyladenosine (m^6^A) modification, or a Kozak sequence(*13–15*). Computational predictions have identified over 328,080 circRNAs with putative coding potential(*16*). However, most lack experimental validation, likely due to low expression levels or inaccurate bioinformatic predictions.

In this study, we explore the possibility of a non-canonical translation mechanism that may generate a vast repertoire of previously unannotated polypeptides. This unconventional mechanism exhibits three fundamental distinctions from canonical translation: (i) initiation by recognition of 5’-GUA-3’ codon (instead of the conventional 5’-AUG-3’), (ii) elongation proceeds in the backward (3’→5’) direction along circular RNA templates, maintaining the triplet codon framework and the universal “genetic code”, but with reversed decoding orientation; and (iii) termination when meeting a distinct set of stop codons 5’-AAU/AGU/GAU-3’ (instead of the conventional 5’-UAG/UGA/UAA-3’) (**Fig. 1A and fig. S1A**). The directionality here refers to the orientation of the sugar-phosphate backbone as conventionally defined in molecular biology. In accordance with the decoding direction, we designate the new translation mode as ‘backward translation’ (BT) to distinguish it from the classical forward (5’→3’) translation (FT) (**Fig. 1A**). Under these definitions, the **backward open reading frame (BW-ORF)** refers to a continuous stretch of codons read in the 3’→5’ orientation on the circular RNA template that can potentially encode a polypeptide. Noteworthy, backward translation is absolutely different from two atypical translation modes: translation of ‘antisense transcript’ and translation of ‘reverse transcript’. In the former, the complementary coding strand, instead of the template strand, is utilized for transcription and subsequent translation (i.e., antisense DNA→mRNA→protein) (**fig. S1B**)(*17*); and in the latter, the mRNA is reversely transcribed into cDNA, which serves as the template for further transcription and translation (i.e., mRNA→cDNA→mRNA→protein)(*18*).

**Fig. 1.**
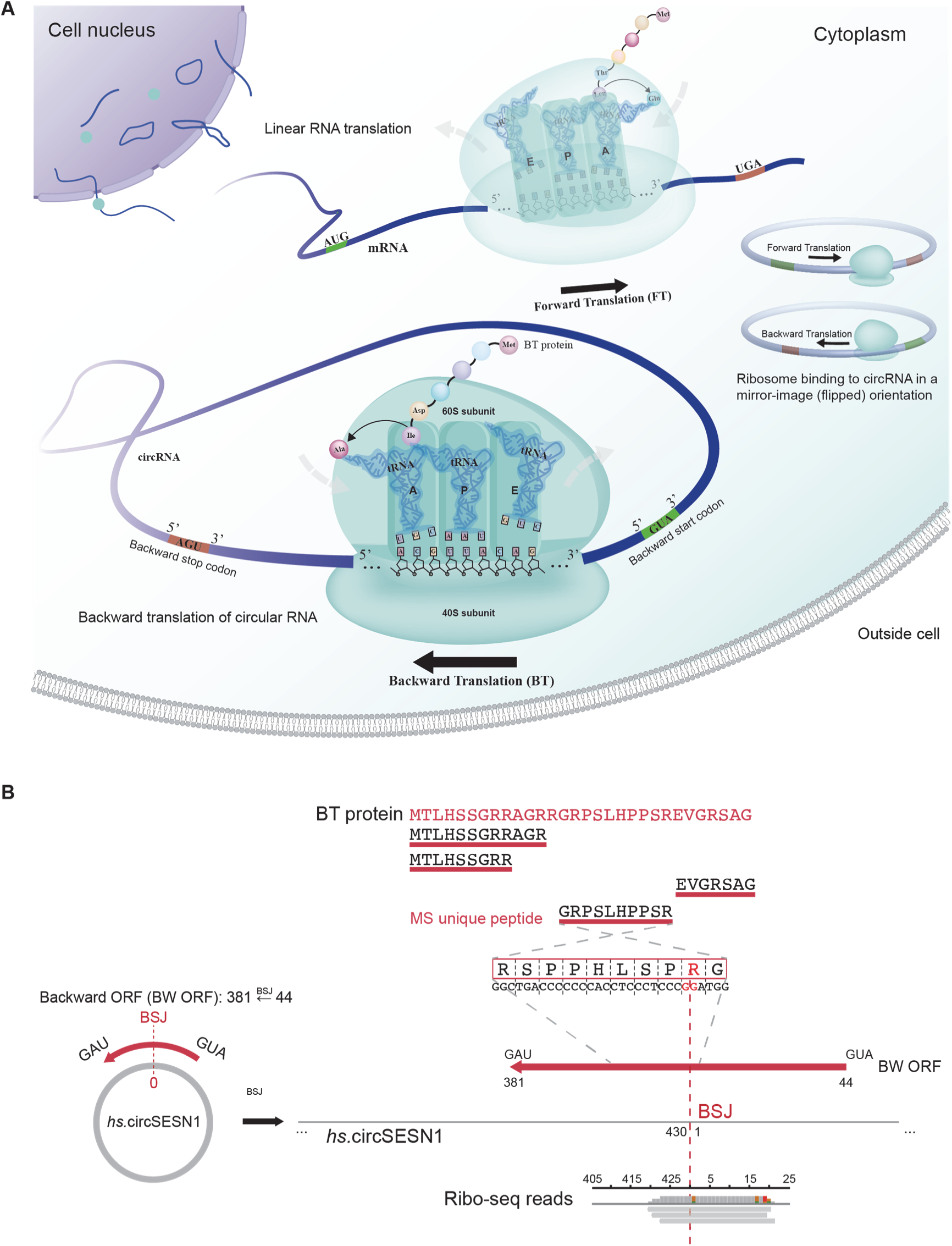
Discovery of circRNA-mediated backward translation (BT). **(A)** Schematic illustration of circRNA-mediated backward translation. The backward translation is initiated by scanning the backward start codon (5’-GUA-3’) and proceeds via decoding triplet genetic codes in the 3’→5’ direction along the circRNA till the stop codons 5’-AAU/AGU/GAU-3’. **(B)** Schematic illustration of the strategy for detecting BT proteins using mass spectrometry and Ribo-seq. BT proteins were validated by identifying at least two unique peptides from MS/MS data that mapped unambiguously to the predicted BT protein sequence. For Ribo-seq support, all reads potentially originating from forward translation or linear transcripts were removed, and the remaining reads were aligned to a 50-nt BT-specific region spanning the BSJ (25 nt upstream + 25 nt downstream). BT translation was considered supported when at least two BSJ-spanning Ribo-seq reads were detected.

### Widespread Presence of CircRNA-mediated Backward Translation

We systematically identified 74,982 circRNAs across 379 transcriptomes from six species (human, mouse, zebrafish, *Drosophila*, nematode, and yeast) with trCirit-BSJ (back-splice junction) annotation **(fig. S2 and Materials and Methods)**, out of 114,221 candidates detected by CIRIT(*19*), and integrated an additional 1,532,085 circRNAs from public databases (**Table S1**). Employing our custom computational pipeline trCIRIT (**fig. S2 Materials and Methods**), we identified 2,674,362 potential BW-ORFs (>50 nt), including 2,264,360 in humans, within 754,496 putatively translatable circRNAs (577,137 human) (**Fig. 2A and Table 1)**.

**Fig. 2.**
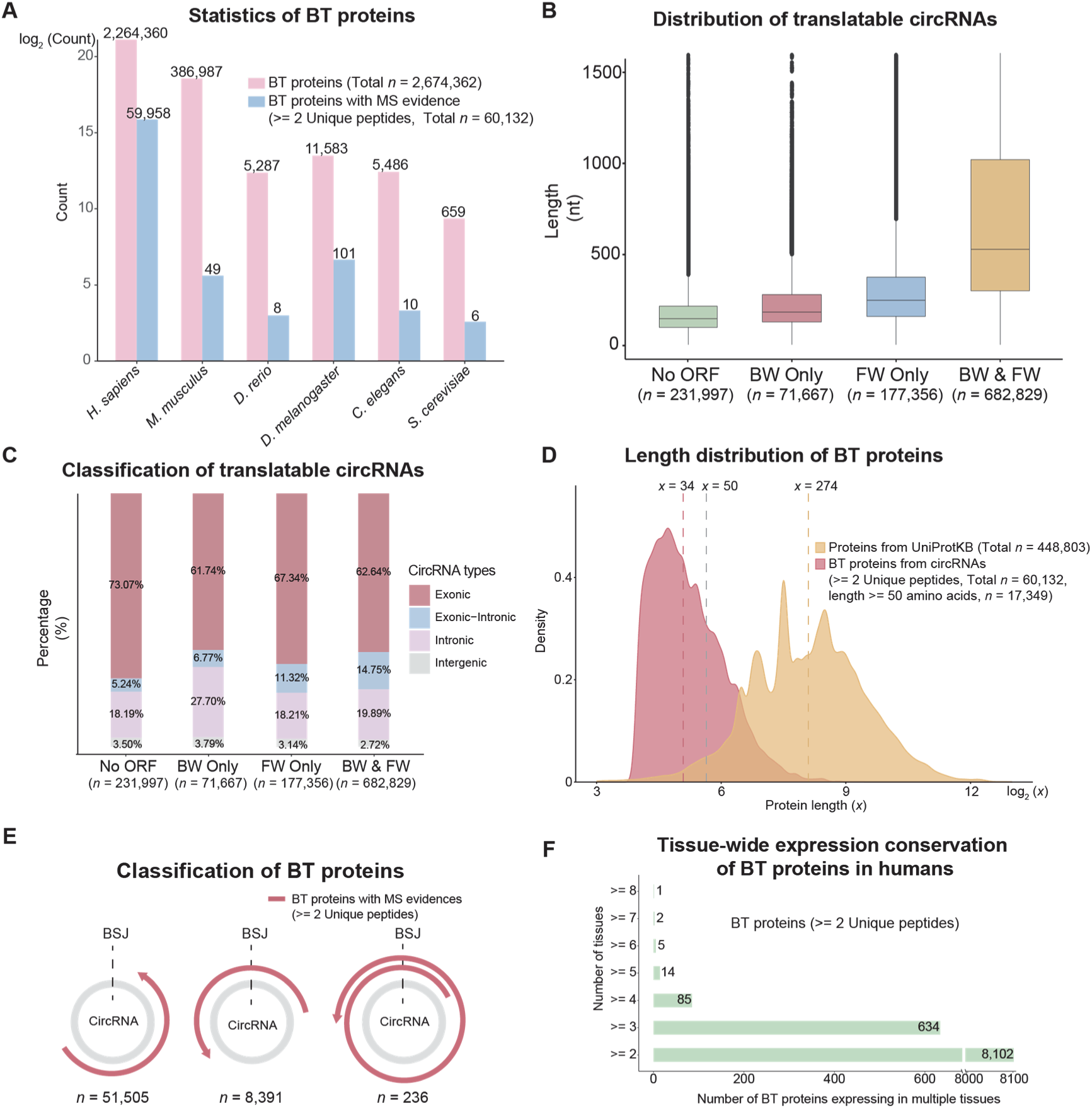
Statistical overview of backward translation. **(A)** Statistics of predicted backward open reading frames (ORFs) across six eukaryotic species: *Homo sapiens*, *Mus musculus*, *Danio rerio*, *Drosophila melanogaster*, *Caenorhabditis elegans*, and *Saccharomyces cerevisiae*. **(B)** Distribution of predicted translatable circRNAs based on sequence length. **(C)** Classification of translatable circRNAs by composition type. **(D)** Length distribution of BT-derived polypeptides/proteins (BT proteins) compared to those encoded by conventional linear mRNAs. **(E)** Classification of BT polypeptides/proteins based on whether translation crosses the back-splice junction (BSJ-crossing). **(F)** Tissue-wide expression conservation of BT polypeptides/proteins in humans. Analysis included 59,863 human BT polypeptides/proteins. Polypeptides were defined as sequences of 10–50 amino acids; proteins, >50 amino acids.

**Table 1.** Statistics of BT circRNAs/BW ORFs supported by MS and/or Ribo-seq evidence.

| Specie | Type | All | MS* | Ribo-seq** |
| --- | --- | --- | --- | --- |
| All | BT circRNAs | 754,496 | 42,591 | 44,930 |
|  | BW ORFs | 2,674,362 | 60,132 |  |
| *Homo sapiens* | BT circRNAs | 577,137 | 42,419 | 23,092 |
|  | BW ORFs | 2,264,360 | 59,958 |  |
| *Mus musculus* | BT circRNAs | 168,236 | 49 | 20,845 |
|  | BW ORFs | 386,987 | 49 |  |
| *Danio rerio* | BT circRNAs | 3,230 | 8 | 155 |
|  | BW ORFs | 5,287 | 8 |  |
| *Drosophila melanogaster* | BT circRNAs | 3,748 | 99 | 488 |
|  | BW ORFs | 11,583 | 101 |  |
| *Caenorhabditis elegans* | BT circRNAs | 1,854 | 10 | 275 |
|  | BW ORFs | 5,486 | 10 |  |
| *Saccharomyces cerevisiae* | BT circRNAs | 291 | 6 | 75 |
|  | BW ORFs | 659 | 6 |  |

\*, supported by $\geq 2$ unique peptides.
\*\*, supported by $\geq 1$ BSJ-spanning reads, excluding contamination of linear mRNA and forward ORF.

Currently, circRNA translation is usually validated through two principal approaches: (i) translation activity inferred from ribosome profiling (Ribo-seq), which captures ribosome-protected footprints on circRNAs, and (ii) direct detection of translated products via mass spectrometry (MS)-based proteomics. Through systematic analysis of 434 matched transcriptomes and Ribo-seq translatomes (**Supplementary Table 2**), we identified active translation of BW-ORFs in 44,930 circular RNAs, including 23,092 human circRNAs (**Supplementary File. 2**). All cases were validated by at least one BW-ORF-specific back-splicing junction (BSJ)-spanning reads after rigorously excluding linear mRNAs, rRNAs, canonical circRNA translation events, and other ncRNAs (**fig. S3**), confirming both their circular nature and translational activity (**Fig. 2B**). Applying more stringent filtering (≥ 2 BSJ-spanning reads) yielded 13,465 high-confidence BT circRNAs supported by Ribo-seq. The failure to map these BSJ-spanning reads to reference genomes argues against sequence artifacts and supports their origin from genuine backward translation. Furthermore, a subset of these BT ORFs exhibited characteristic hallmarks of canonical translation—including in-frame read enrichment and reduced signal at start/stop sites—despite their overall low read coverage. (**fig. S5**). However, given the inherent limitations of Ribo-seq—including its inability to discriminate between active translation and ribosomal scanning, coupled with low sensitivity for rare or short ORFs—we established MS as the definitive benchmark for BT validation.

Proteomic validation was performed through a large-scale reanalysis of 6,112 public MS/MS datasets in accordance with HUPO guidelines (**Supplementary Table 1**). Using standard filtering criteria (≥1 unique peptide, FDR <1%), we identified 294,079 BT-derived polypeptides (hereafter BT proteins; **fig. S4 and Materials and Methods**). Applying more stringent criteria (≥2 unique peptides, FDR <1%) yielded 60,132 high-confidence BT proteins (**Fig. 2A and Supplementary File. 3**), of which 59,958 were of human origin—a set that nearly triples the number of entries in the current curated human proteome (UniProtKB: 20,421). Notably, 25 of these high-confidence proteins were further supported by orthogonal evidence (≥2 BSJ-spanning Ribo-seq reads and ≥1 BSJ-spanning unique peptide; **Fig. 1B**). This unprecedented expansion of the known proteome fundamentally reshapes our understanding of circRNA biology, establishing backward translation as a major contributor to proteomic diversity.

Statistical analyses reveal that longer circRNAs (>500 nucleotide, nt) exhibit greater translational potential and a higher propensity for bidirectional translation compared to shorter circRNAs (**Fig. 2B**). Among circRNA types, exon-containing circRNAs dominate predicted BT events (>75%), while only ∼20% of intronic circRNAs retain BT capability—a ratio similar to that observed in forward translation (FT) (**Fig. 2C**). Notably, approximately 69% of BT-derived proteins are small polypeptides (20-60 amino acids, aa), averaging one-eighth the length of canonical linear mRNA-encoded proteins (UniProtKB) (**Fig. 2D**). The largest MS-validated BT protein (695 aa) identified in this study was encoded by yeast *sc.*circFLO11 (1,042nt) in a rolling-circle translation mode. Based on translation patterns, 51,505 BT proteins do not span BSJs, 8,391 cross BSJs once, and 236 predict rolling-circle translation by multiple BSJ crossings (**Fig. 2E**).

CircRNA-mediated backward translation was detected across a broad evolutionary spectrum, from unicellular yeast to complex multicellular organisms, including mouse and human (**Fig. 2A**). While present in diverse species, the majority of BT proteins identified in this study derive from human samples—a reflection of the relative depth and availability of human multi-omics datasets. Within humans, BT proteins were found in multiple tissues, with their detection rate closely linked to the coverage and quality of the underlying proteomic data. Among the 59,863 high-confidence human BT proteins, 51,761 (≈86.5%) appear to be tissue-restricted, whereas 8,102 (approximately 13.5%) were observed in two or more tissues (**Fig. 2F**).

### Sequential, Expression, and Structural Features of BT Proteins

Unlike conventional circRNA-derived translation products (i.e., the FT proteins), which frequently encode truncated variants of their host proteins (often featuring altered C-terminal sequences or tandem domain repeats) (*10*), BT proteins exhibit entirely distinct sequence features that arise from their unique 3’→5’ decoding mechanism, fundamentally distinguishing them from the products of their cognate linear mRNAs. Furthermore, our comprehensive homology analysis revealed no significant matches to known proteins when aligning BT proteins against the UniProtKB database and PeptideAtlas database(*20*) (2,531,100 annotated human peptides) using stringent criteria (>80% sequence identity and coverage) (**Supplementary Table 3-1**). We further mined potential functional domains within BT proteins using the SMART server(*21*). Despite screening for mobile element-derived ORFs and known domain architectures, no conserved domains were identified by SMART. In contrast, a systematic search of the PROSITE database(*22*) identified 87 putative motifs, present in ∼80% of BT proteins (**Supplementary Table 3-2**). The ten most prevalent motifs accounted for 80% of all detected PROSITE signatures, involved in several essential cellular functions, including metabolism, signal transduction, and post-translational modifications.

To determine whether BT protein abundance is regulated at the level of circRNA availability or canonical translation, we analyzed a paired transcriptome-proteome dataset of 70 human liver samples (PXD006512) (*23*). We found that BT proteins were on average ∼32-fold less abundant than their cognate linear mRNA-encoded host proteins (**fig. S6A**). Nevertheless, a subset of 2,502 BT proteins reached expression levels comparable to those of canonical proteins, supporting their potential biological relevance. Notably, neither circRNA expression nor host protein abundance showed a meaningful correlation with BT protein levels (circRNA vs. BT protein: *R*² = 3*e*⁻⁵; host protein vs. BT protein: *R*² = 2*e*⁻³; **fig. S6B and C**). These results indicate that BT protein output is not simply a passive reflection of circRNA availability or host gene translation, but may be subject to dedicated regulatory control.

Moreover, we selected five BT proteins with diverse lengths and predicted their structures using AlphaFold (**fig. S6D**). The simulation demonstrated that these BT proteins exhibit pronounced loop-rich regions and high conformation flexibility. Further structural alignment of these BT proteins against 542,337 AlphaFold-predicted protein models via K-means clustering revealed minimal structural similarity to known proteins (**fig. S6D**).

### Endogenous Validation of Backward Translation

To experimentally validate endogenous backward translation, we selected human circCAPN15 as a model circRNA, which was computationally predicted to encode a 234-amino acid BT protein (*hs*.circCAPN15_*bt*_*p*; 25.8 kDa). In the human genome, *hs.*circCAPN15 resides in the intronic region of *CAPN15* (a known regulator of cell proliferation, growth, and migration(*24*)). Divergent primer-based RT-PCR (**Supplementary Table 4**) confirmed *hs.*circCAPN15 expression in HeLa cells, RNase R resistance validated its circular nature, while Sanger sequencing confirmed the BSJ (**fig. S7A**). Immunofluorescence using two specific antibodies (anti-IgG as a negative control) detected cytoplasmic localization of the BT protein (**Fig. 3A**), confirming the endogenous backward translation of *hs.*circCAPN15. The specificity of the customized antibodies was further evaluated by gene knockout experiments shown below.

**Fig. 3.**
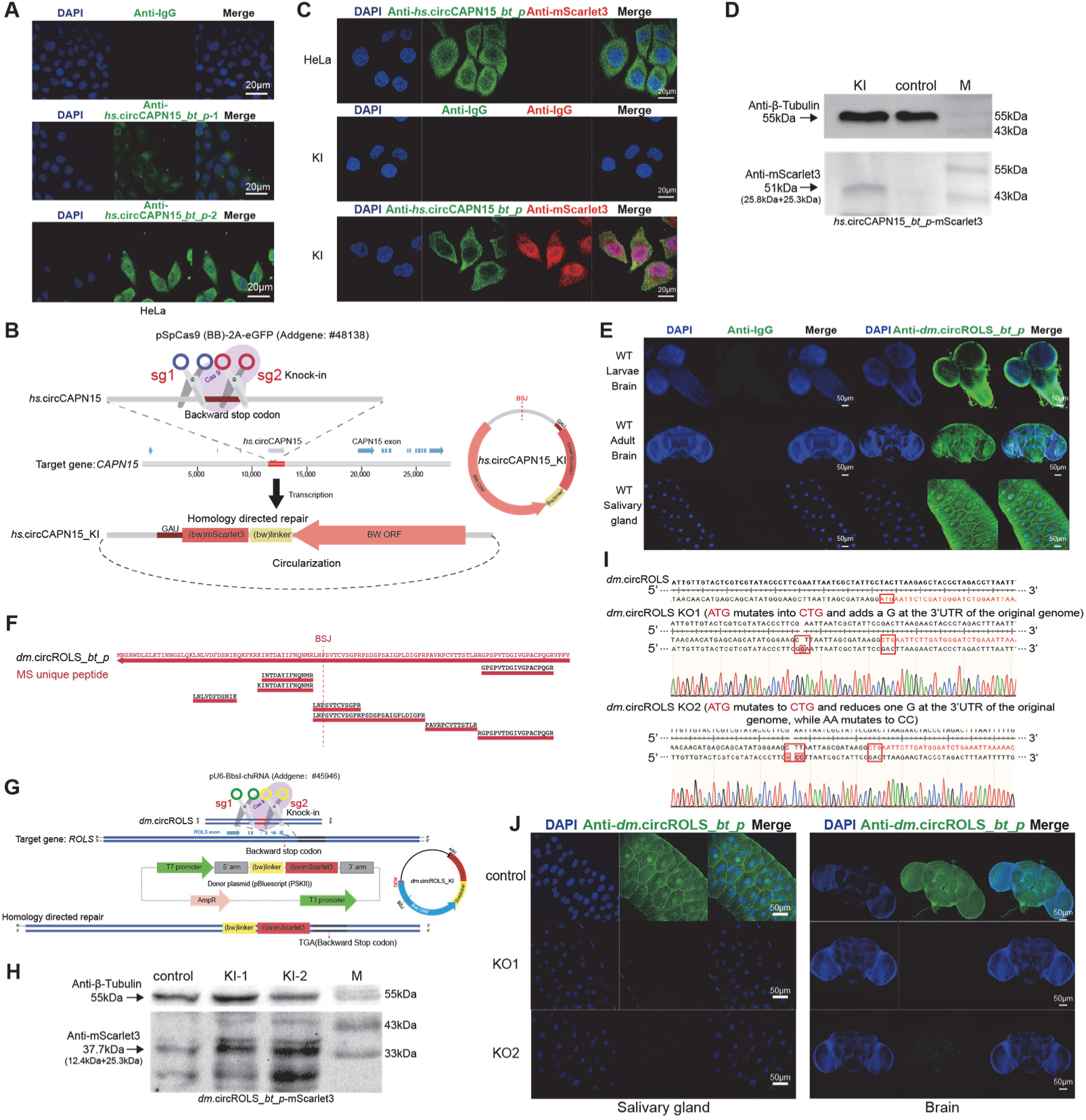
*In vitro* and *in vivo* validation of endogenous backward translation. **(A)** Immunofluorescence validation of endogenous backward translation of *hs.*circCAPN15 in HeLa cells using custom-made antibodies. Two anti-*hs.*circCAPN15*_bt_p* antibodies were used (green signal), with IgG as the negative control. Nuclei were counterstained with DAPI (blue). Representative confocal images are shown (Scale bar, 20 μm). **(B)** Schematic of the plasmid design for the knock-in of *hs.*circCAPN15. A reversed open reading frame (ORF) encoding the red fluorescent protein mScarlet3 was inserted into the genomic region corresponding to the BW-ORF of *hs*.circCAPN15, immediately upstream of its stop codon (AGT). The knock-in was achieved using CRISPR/Cas9 technology. **(C)** Confocal images showing successful knock-in of *hs.*circCAPN15 to express the backward-translated fluorescent fusion protein (*hs.*circCAPN15*_bt_p*-mScarlet3) in HeLa cells. mScarlet3 fluorescence (red) indicates successful knock-in and colocalizes with anti-*hs.*circCAPN15*_bt_p* antibody staining (green). DAPI marks nuclei (blue). Scale bar, 20 μm. **(D)** Western blot (WB) analysis using an anti-mScarlet3 antibody to detect the BT fusion protein in wild-type control and *hs*.circCAPN15*_bt_p*-mScarlet3 knock-in (KI) HeLa cells. The arrow marks the expected *hs.*circCAPN15*_bt_p*-mScarlet3-BT fusion protein band (51.1 kDa; *hs.*circCAPN15*_bt_p*: 25.8 kDa, mScarlet3: 25.3 kDa). β-Tubulin (55 kDa) was used as a loading control. **(E)** Endogenous backward translation of *dm.*circROLS*_bt_p* in *Drosophila*. Representative confocal images of larval and adult brains, as well as larval salivary glands, from wild-type (WT) *Drosophila* stained with anti-*dm.*circROLS*_bt_p* antibody (green). IgG served as a negative control. DAPI was used for nuclear counterstaining (blue). Scale bar, 50 μm. **(F)** Schematic illustration of unique peptides of *dm*.circROLS*_bt_p* detected by IP–MS (DIA). **(G)** Schematic of the plasmid design for the knock-in of *dm.*circROLS. A reversed mScarlet3 ORF was inserted just before the stop codon (AGT) in the backward ORF of *dm.*circROLS. CRISPR/Cas9 was used for knock-in. **(H)** Western blot analysis of mScarlet3 expression in wild-type control and two knock-in (KI-1 and KI-2) *dm.*circROLS-mScarlet3 *Drosophila* lines. The expected BT fusion protein band (37.7 kDa; *dm.*circROLS*_bt_p*: 12.4 kDa, mScarlet3: 25.3 kDa) is indicated by an arrow. β-Tubulin (55 kDa) served as a loading control. Additional bands represent nonspecific signals. **(I)** Sanger sequencing confirmed successful knockout of *dm.*circROLS*_bt_p*. Two mutant alleles (KO-1 and KO-2) were generated via CRISPR/Cas9. Both mutations altered the backward start codon (3’-ATG-5’ to 3’-CTG-5’) and introduced a premature backward stop codon after the third amino acid, through synonymous mutations that preserved the wild-type coding sequence of the host *ROLS* gene but abolished backward translation. **(J)** Immunofluorescence detection of the translated *dm.*circROLS*_bt_p* product in the adult brains and salivary glands of third-instar larvae from wild-type control, KO-1, and KO-2 *Drosophila*. Green fluorescence indicates antibody staining for *dm*.circROLS*_bt_p*. Nuclei were counterstained with DAPI (blue). Representative confocal images are shown. Scale bar, 50 μm.

Using CRISPR/Cas9, we inserted the reversed mScarlet3 ORF into the *CAPN15* locus, placing it upstream of the stop codon (5’-AGT-3’) in *hs*.circCAPN15 backward ORF (**Fig. 3B**). This knock-in strategy enabled HeLa cells transcription of engineered *hs*.circCAPN15 encoding the backward translated *hs.*circCAPN15*_bt_p*-mScarlet3 fusion protein (∼51kDa). Correct knock-in was confirmed by Sanger sequencing (**fig. S7B**). Immunofluorescence revealed BT protein signals of reverse-coded mScarlet3 and *hs.*circCAPN15 exclusively in the engineered cells, absent in the wild-type cells, demonstrating backward translation of *hs.*circCAPN15 (**Fig. 3C**). The conclusion was further supported by Western blot analysis of mScarlet3 fusion BT protein (**Fig. 3D**).

Endogenous BT expression was further validated in *Drosophila dm*.circROLS (see also below for functional studies), an exonic circRNA transcribed from the complementary strand of the *ROLS* gene according to our computational analysis, which likely encodes a 113-aa BT protein. RNase R-resistant RT-PCR confirmed its circularity, and Sanger sequencing verified the BSJ (**fig. S7C**). Immunofluorescence staining using target-specific antibodies revealed strong *dm*.circROLS-derived BT protein expression in both brain and salivary gland tissues (**Fig. 3E**), providing *in vivo* evidence of endogenous backward translation. The additional immunoprecipitation assay successfully captured *dm.*circROLS*_bt_p*-derived signals, which were subsequently identified by MS spectrometry (**Fig. 3F and Supplementary Table 1-3**). In the same way, we utilized CRISPR/Cas9 to generate a knock-in line, inserting a reversed mScarlet3 ORF in-frame preceding the backward stop codon of *dm.*circROLS (5’-AGT-3’; **Fig. 3G**). The integration was verified by Sanger sequencing (**fig. S7D**). Western blot confirmed the production of *dm.*circROLS*_bt_p*-mScarlet3 fusion protein (∼37.7kDa) (**Fig. 3H**). Moreover, we created two knockout lines by mutating the backward start codon (5’-GUA-3’ to 5’-GUC-3’) of *dm.*circROLS while preserving the coding capability of host gene *ROLS* (**Fig. 3I**). Immunofluorescence revealed markedly reduced BT protein signals of *dm.*circROLS in the brain and salivary gland tissues of the two knockout lines (**Fig. 3J**), confirming successful ablation of backward translation.

Together, both human cells and *Drosophila* studies verified the endogenous backward translation in eukaryotic systems.

### *In vitro* Engineering of Backward Translation

Beyond their natural occurrence, BT proteins can be engineered for expression using synthetic biology approaches. Taking a commercially available plasmid (pcDNA3.1+), previously validated for circRNA-dependent forward translation(*25*), we designed a BT protein synthesis system by incorporating two split reverse eGFP (enhanced green fluorescence protein) ORF segments and the EMCV-IRES element in the plasmid construct, namely circ-EMCV-IRES-(bw)-eGFP (**Fig. 4A**). The green fluorescence signal of eGFP could be detected only if the split reverse eGFP ORF segments are correctly circularized and expressed via backward translation (**Fig. 4B**). As expected, HEK-293T cells transfected with this engineered circ-EMCV-IRES-(bw)-eGFP construct expressed circRNAs (**Fig. 4C**, measured with qPCR) and underwent backward translation (**Fig. 4D**).

**Fig. 4.**
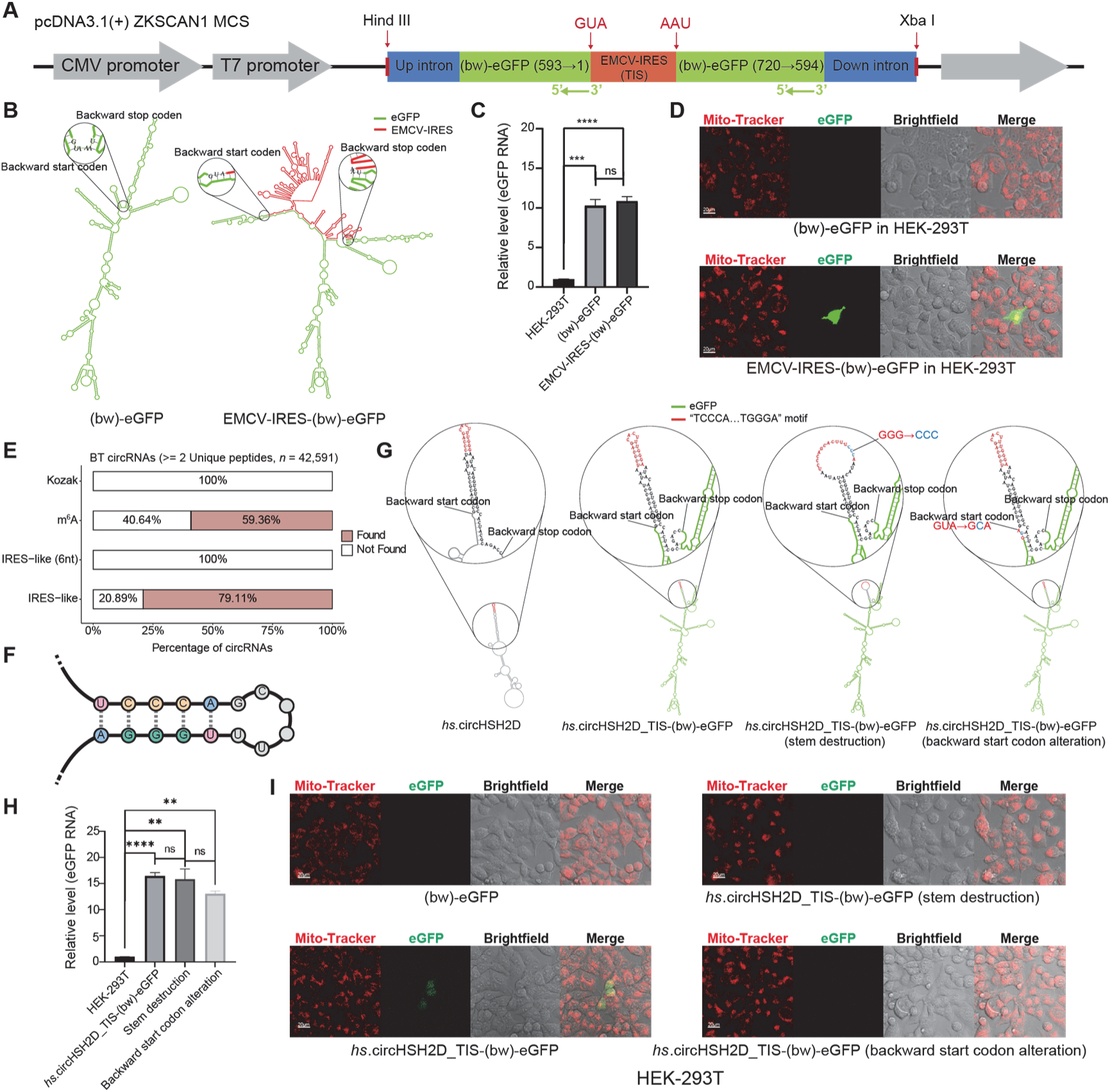
Sequence characteristics associated with the initiation of backward translation. **(A)** Schematic of vector design for backward translation of enhanced green fluorescent protein (eGFP) in human cells. The pcDNA3.1 vector was used as the backbone for circRNA expression. The construct consisted of two reverse eGFP fragments (coordinates: 593→1 and 720→594), separated by a translation initiation site (TIS)-containing sequence. The TIS sequence was flanked by a backward start codon (←GUA) and a backward stop codon (AAU←). TIS elements used in this study included either upstream sequences derived from endogenous backward ORFs or the EMCV internal ribosome entry site (IRES). **(B)** Structural illustration of two circ-(bw)-eGFP constructs: one lacking a TIS sequence element and one incorporating EMCV-IRES as TIS. The eGFP sequence is shown in green, and the TIS region is marked in red. **(C)** RT-qPCR quantification of reverse eGFP expression in HEK-293T cells transfected with empty vector, reverse eGFP without TIS, or reverse eGFP containing EMCV-IRES as TIS (*n*=3). Tubulin was used as the internal reference gene. Primers spanning the back-splice junction (BSJ) were used to specifically detect circular RNA. All RNA samples were treated with RNase R to enrich for circular species. **(D)** Representative confocal microscopy images showing eGFP fluorescence (green) resulting from backward translation of circ-(bw)-eGFP in HEK-293T cells. Fluorescence was observed in cells transfected with reverse eGFP containing EMCV-IRES, but not in those lacking TIS. Scale bar, 20 μm. The fluorescent signal was detected in ∼1–2 out of every 100,000 cells. Live cells were stained with Mito-Tracker (red, mitochondria). **(E)** Summary of four sequence features known to promote circRNA translation upstream of backward ORFs: IRES-like elements predicted by IRESfinder, IRES-like hexamers aligned to the 10-nt region upstream of the backward start codon, Kozak consensus sequences identified by in-house scripts, and N6-methyladenosine (m⁶A) sites predicted using SRAMP. **(F)** The secondary structure of conserved motif ‘TCCCA…TGGGA’ identified from backward-translated circRNAs using MEME analysis. **(G)** Predicted secondary structures of circRNAs involved in regulating backward translation, modeled using RNAfold. From left to right: *hs.*circHSH2D; circ-(bw)-eGFP incorporating *hs.*circHSH2D-derived TIS; circ-(bw)-eGFP with the same TIS but with a stem-loop structure disrupted by mutation; and circ-(bw)-eGFP in which the backward start codon (5’-GUA-3’) was mutated to 5’-GCA-3’. The latter two constructs were predicted to be untranslatable. **(H)** RT-qPCR quantification of reverse eGFP in HEK-293T cells transfected with empty vector, *hs.*circHSH2D-TIS-(bw)-eGFP, *hs.*circHSH2D-TIS-(bw)-eGFP with stem-loop disruption, and *hs.*circHSH2D-TIS-(bw)-eGFP with start codon mutation (n=3). Tubulin was used as the internal control. Primers spanned the BSJ, and RNase R treatment was applied to enrich for circRNAs. **(I)** Representative confocal microscopy images showing eGFP fluorescence (green) from backward translation of circ-(bw)-eGFP in HEK-293T cells transfected with constructs listed in panel h. Fluorescence was observed only in cells transfected with *hs.*circHSH2D-TIS-(bw)-eGFP. No signal was detected in other groups. Scale bar, 20 μm. live cells were stained with Mito-Tracker (red, mitochondria). Statistical comparisons of eGFP expression levels under different conditions were performed using Mann-Whitney U in GraphPad Prism (version 9.2.0). Statistical significance: *ns*: not significant, “*”: *p* <0.05, “**”: *p* <0.01, “***”: *p* <0.001, “****”: *p* <0.0001.

As circ-(bw)-eGFP without IRES could not undergo backward translation (**Fig. 4D**), we reasoned that circRNAs may require specialized initiation elements for initiating translation. Potential candidates included IRES-like sequences, 6nt IRES-like sequences, Kozak sequence, as well as m^6^A modification that were previously identified for circRNA forward translation (*9*). Using IRESfinder software v.1.1.0(*^26^*), we analyzed upstream sequences of high-confidence BW-ORFs (MS-supported with ≥2 unique peptides) from 42,591 BT circRNAs, identifying IRES-like motifs in 79.11% of cases (**Fig. 4E and Supplementary Table 5**). None of them contained 6-nt IRES-like motifs or Kozak sequences (reversed), suggesting these elements are dispensable for backward initiation. 59.36% of BT circRNAs exhibited predicted m^6^A modifications (**Fig. 4E and Supplementary Table 5**). Notably, our motif enrichment analysis using MEME v5.3.0^22^ uncovered 10 conserved sequence motifs in human BT circRNAs that appear to mediate backward translation initiation (**Supplementary Table 6-1**). Of them, the most abundant motif, ‘TCCCA…TGGGA’ (present in two conserved motifs), can form a stable stem-loop structure and appears to function as a core element of backward translation initiation sites (TISs) (**Fig. 4F**). Additional motif search in all studied BT circRNAs with MAST v4.11.2 software discovered that approximately 22.94% of cases contained this motif (**Supplementary Table 6-2**). It is noteworthy that prior to this study, prevailing research has generally held that eukaryotic IRES elements exhibit little conservation in either primary sequence or secondary RNA structure(*27*).

To validate our analysis, we extracted three distinct putative TIS sequences from *hs*.circHSH2D, *hs*.circITIH4, and *hs*.circMGC27382, utilizing them to initiate backward translation of circ-(bw)-eGFP. Secondary structure simulation of circ-(bw)-eGFP showed that these three TISs formed stem-loop structures preceding the (bw)-eGFP ORF as predicted with RNAfold (**Fig. 4G and fig. S8A**). Notably, for the TISs derived from *hs*.circITIH4 and *hs*.circMGC27382, the “TCCCA…TGGGA” motif did not predict to be involved in forming of stem-loop structure directly after insertion into the circ-(bw)-eGFPs. However, transfection of these plasmid constructs into HEK-293T cells all successfully transcribed circRNAs (**Fig. 4H**) and elicited green fluorescence (with Mito-tracker as a mitochondrial probe in living cells), indicating the backward translation of eGFP (**Fig. 4I and fig. S8B**). Mutation experiments that either disrupted the stem structure of the ‘TCCCA…TGGGA’ motif or altered the backward start codon (5’-GUA-3’→5’-GCA-3’) could interrupt the backward translation of circ-(bw)-eGFP (**Fig. 4G to I**). In addition, we extracted two TISs from *hs*.circCAPN15 and one TIS from *hs*.circIFITM1, all of which were predicted to form typical stem-loop structures but lacked the ‘TCCCA…TGGGA’ motif (**fig. S8C**). The placement of these TISs before the (bw)-eGFP ORF all successfully drove the backward translation (**fig. S8D**). Notably, two TISs from different regions of *hs.*circCAPN15 were able to initiate BT, indicating the presence of multiple TISs within a single circRNA. We further explored the conservation of these TISs across species, extracting two TISs from *Drosophila*, specifically *dm*.circSCRIB and *dm*.circUNC-5 (**fig. S8E**), both of which enabled the backward translation of circ-(bw)-eGFP in HEK-293T cells (**fig. S8F**).

To eliminate the possibility of false-positive BT results caused by aberrant plasmid expression, we synthesized two circRNAs, *hs.*circIFITM1-TIS-(bw)-eGFP and *hs.*circDENND5B-TIS-(bw)-eGFP, *in vitro*. The circularization and correctness of synthetic circRNAs were confirmed by RT-PCR (**Fig. 5A**) and Sanger sequencing (**Fig. 5B**), respectively. Transfection of either synthetic circRNAs into HEK-293T cells elicited robust green fluorescence in a subset of cells (with Mito-tracker as a mitochondrial probe in living cells), confirming successful backward translation of eGFP from the engineering circular RNA templates (**Fig. 5C**). The exogenous BT products (eGFP, 27kDa) from the circ-(bw)-eGFPs were further confirmed by immunoprecipitation (IP) analysis with a specific antibody against eGFP (**Fig. 5D**).

**Fig. 5.**
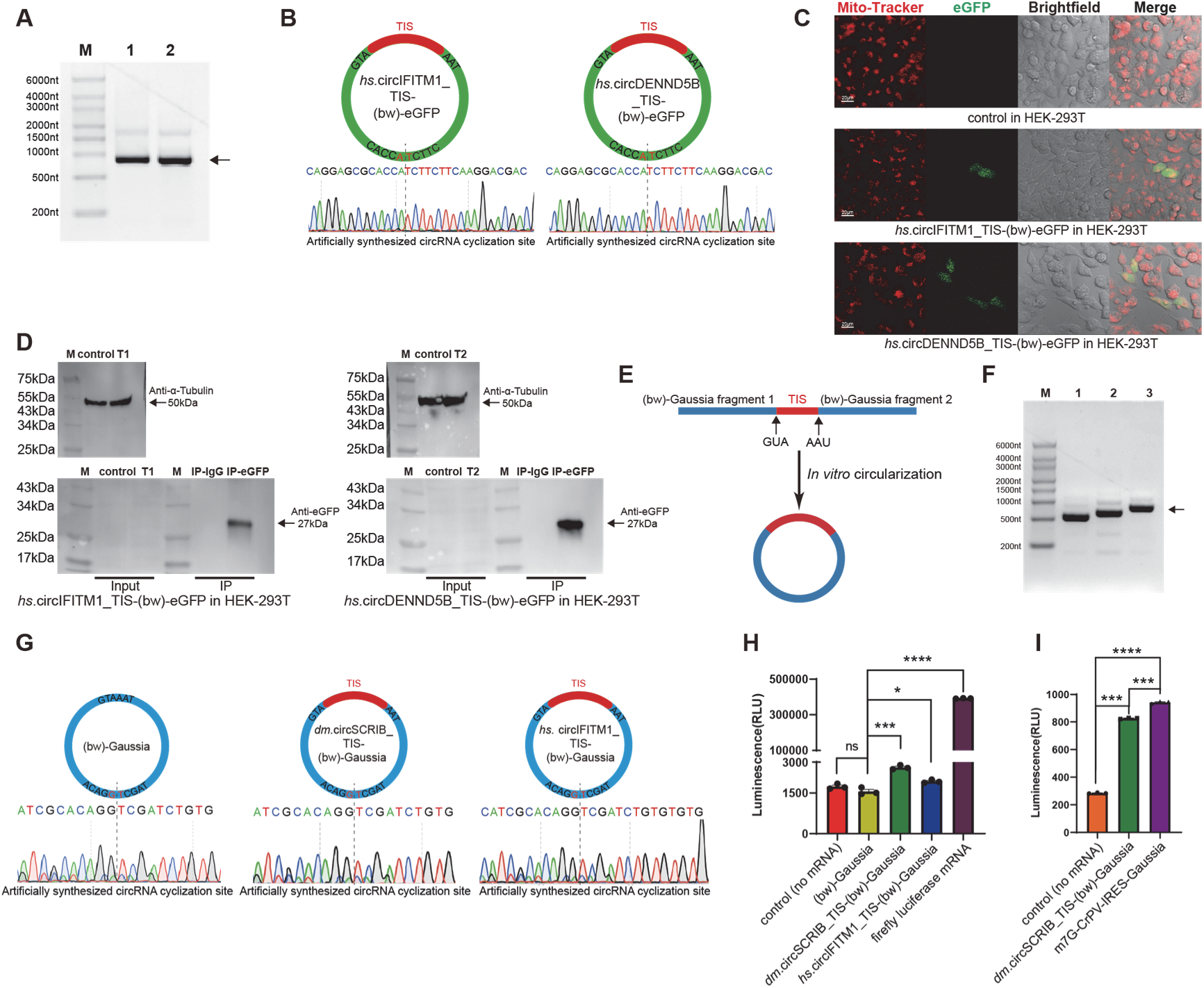
*In vitro* translation of backward ORFs in circular RNAs. **(A)** Synthetic circular RNAs, *hs.*circIFITM1-TIS-(bw)-eGFP (819 nt, Lane 1) and *hs.*circDENND5B-TIS-(bw)-eGFP (831 nt, Lane 2), were analyzed by electrophoresis on a 2% agarose gel (125 V, 25 min). **(B)** Sanger sequencing of RT-PCR products from *hs*.circIFITM1-TIS-(bw)-eGFP and *hs.*circDENND5B-TIS-(bw)-eGFP, reverse-transcribed with random primers and amplified using back splice-junction-specific primers. **(C)** Electroporation of synthetic *hs.*circIFITM1-TIS-(bw)-eGFP and *hs.*circDENND5B-TIS-(bw)-eGFP into HEK-293T cells. After 24 h, live cells were stained with Mito-Tracker (red, mitochondria). The eGFP fluorescence (green) indicated expression from the circRNA constructs. Scale bar, 20 μm. **(D)** Immunoblot analysis of whole-cell lysates (input) and eGFP immunoprecipitates (IP) from HEK-293T cells transfected with either *hs.*circIFITM1-TIS-(bw)-eGFP (T1) or *hs.*circDENND5B-TIS-(bw)-eGFP (T2), alongside untransfected controls. The eGFP was detected using an anti-eGFP antibody, with a predicted band at 27 kDa. The α-Tubulin (50 kDa) was used as a loading control. **(E)** Schematic representation of the synthetic circGaussia reporter used in cell-free translation. **(F)** Electrophoresis of *in vitro*–synthesized circular RNAs (bw)-Gaussia (558 nt, Lane 1), *hs.*circIFITM1-TIS-(bw)-Gaussia (657 nt, Lane 2), and *dm.*circSCRIB-TIS-(bw)-Gaussia (782 nt, Lane 3) on a 2% agarose gel (125 V, 25 min). **(G)** RT-PCR amplification of the back-splice junctions of *hs.*circIFITM1-TIS-(bw)-Gaussia, (bw)-Gaussia, and *dm.*circSCRIB-TIS-(bw)-Gaussia using divergent primers. Sanger sequencing confirmed correct circularization. **(H)** *In vitro* translation of synthetic *hs.*circIFITM1-TIS-(bw)-Gaussia and *dm.*circSCRIB-TIS-(bw)-Gaussia RNAs using rabbit reticulocyte lysate (*n* =3). Translation was measured by chemiluminescence from Gaussia luciferase activity (coelenterazine substrate, λ =480 nm). Negative Control (no mRNA): RNA-free in vitro translation system; positive control: linear firefly luciferase mRNA. (bw)-Gaussia: Represents the Gaussia circular RNA without any TIS. **(I)** The translation of *dm*.circSCRIB TIS-(bw)-Gaussia RNA (*n*=3), synthesized using the *Drosophila* ovarian extract in vitro translation system, was measured by chemiluminescence detection of Gaussia luciferase activity using the substrate coelenterazine (λ = 480 nm). The positive control consisted of a linear, capped (m7G) Gaussia mRNA containing the Cricket paralysis virus internal ribosome entry site (CrPV IRES). Statistical comparisons between groups were performed using Mann-Whitney U in GraphPad Prism (version 9.2.0). Statistical significance: *ns*: not significant, “*”: *p* <0.05, “**”: *p* <0.01, “***”: *p* <0.001, “****”: *p* <0.0001.

To further eliminate potential false positives of BT caused by other unknown cellular transcription events, we attempted to validate synthetic circRNAs BT in an *in vitro* cell-free translation system (**Fig. 5E**). We synthesized three specific circRNAs, *hs.*circIFITM1-TIS-(bw)-Gaussia and *dm.*circSCRIB-TIS-(bw)-Gaussia, each harboring a reverse ORF encoding the bioluminescent protein Gaussia with the indicated TIS, and circ-(bw)-Gaussia without any TIS as a control. The circularization and correctness of synthetic circRNAs were confirmed by RT-PCR (**Fig. 5F**) and Sanger sequencing (**Fig. 5G**), respectively. To determine whether circRNAs can serve as templates for backward translation, we employed two independent cell-free systems: nuclease-treated rabbit reticulocyte lysate (RRL) and *Drosophila* ovarian extract. In RRL assays, *dm*.circSCRIB-TIS-(bw)-Gaussia generated a 40.7% luminescence increase relative to the no-RNA control, while *hs*.circIFITM1-TIS-(bw)-Gaussia produced an 11.0% increase, indicating detectable BT activity (**Fig. 5H**). As expected, linear firefly luciferase mRNA elicited strong luminescence, confirming efficient conventional translation. In contrast, a TIS-deficient construct yielded no measurable signal, consistent with negative controls.

Using the *Drosophila* extract system, we further tested BT activity. Here, *dm*.circSCRIB-TIS-(bw)-Gaussia showed a 190.02% increase in luminescence over the control, whereas the circRNA-based forward-translation control (m⁷G-CrPV-IRES-Gaussia) reached 231.03% (**Fig. 5I**). Notably, the luminescence magnitudes of forward- and backward-translating circRNAs were comparable, yet both remained substantially lower than that of the linear mRNA positive control. The consistent signal output from BT-competent constructs—and its absence in TIS-deficient counterparts— demonstrates that BT is not a stochastic artifact but a template-directed process amenable to sequence-specific regulation.

Together, these controlled experiments establish backward translation as a genuine process, distinct from transcriptional artifacts. They further imply a regulatory role for circRNA sequence or structure, and showcase how synthetic biology can directly engineer BT—a capability that enables future exploration of its biological significance and practical applications.

### Evidence of Biological Functions of Backward-translated Proteins

Having established the widespread occurrence of circRNA backward translation (BT), we next sought to examine its functional relevance by targeting the BT-derived protein coding sequence of *hs*.circCAPN15 using CRISPR/Cas9. This circRNA originates from an intronic region of the *CAPN15* gene and does not interact with the major *CAPN15* transcript, although it partially overlaps with some rare isoforms. To disrupt the production of the *hs*.circCAPN15 BT protein (234 aa, 25.8 kDa), we generated three independent knockout (KO) HeLa cell lines (*hs*.circCAPN15_*bt_p* KO; **Fig. 6A**). These lines carried either a translation start codon deletion or frame-shift mutations leading to protein truncation, as validated in **fig. S9A**. Immunofluorescence analysis confirmed complete ablation of the predicted protein product (**Fig. 6B**). Notably, qPCR analysis revealed only marginal changes in circRNA levels that did not reach statistical significance (*p* >0.05) (**fig. S9B**). Crucially, these genetic manipulations specifically affected the circRNA-derived BT protein without significantly altering *CAPN15* expression at either transcriptional or translational levels (**Fig. 6C and fig. S9C**). Following the knockout of *hs.*circCAPN15_*bt_p*, the growth of HeLa cells was significantly (average 18.92%) higher than that of the control group (**Fig. 6D and fig. S9D**). In wound healing assays, all three KO cell lines exhibited comparatively better (average 48.7% enhancement) healing capability at 72 hours, indicating that the absence of *hs.*circCAPN15_*bt_p* may promote cell migration (**Fig. 6E and fig. S9E**). Conversely, overexpression of *hs*.circCAPN15*_bt_p* from a linear RNA (the corresponding BW-ORF) template inhibited HeLa cell proliferation and reduced their wound-healing capability (**Fig. 6E**). Note that this overexpression led to an increased abundance of *hs.*circCAPN15_*bt_p* in the cells (**fig. S9F**) without significantly altering the levels of *CAPN15* host protein (**Fig. 6C**). These results clearly indicated that the BT protein encoded by *hs.*circCAPN15 can suppress cellular growth and migration *in vitro*.

**Fig. 6.**
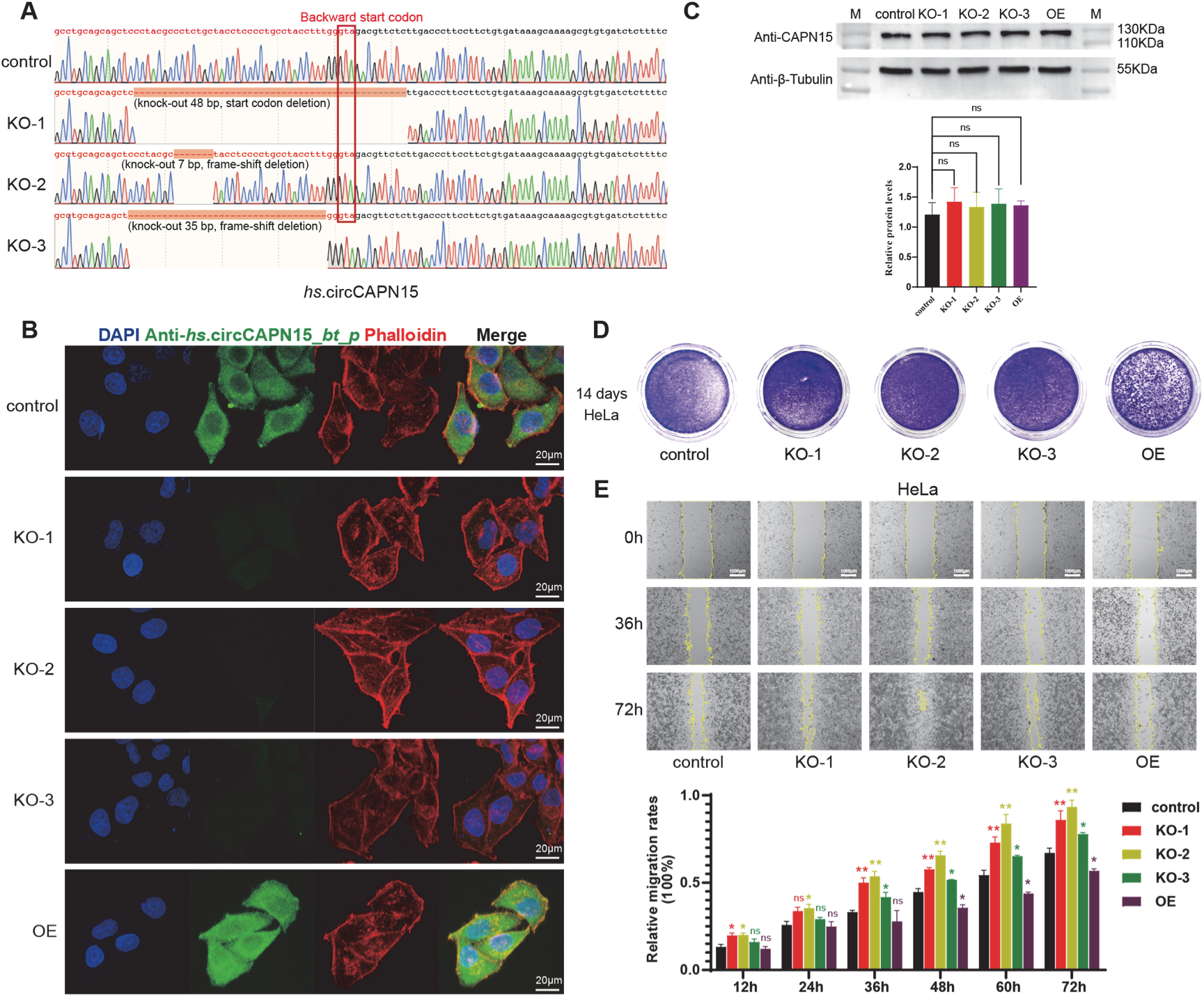
Biological functions of *hs*.circCAPN15*_bt_p* in HeLa cells. **(A)** Sanger sequencing validation of three independent CRISPR/Cas9-mediated knockouts (KOs) of *hs.*circCAPN15*_bt_p*, including a start codon mutation (KO-1) and two frameshift mutations (KO-2 and KO-3). **(B)** Immunofluorescence analysis confirming successful knockout of *hs.*circCAPN15*_bt_p*. Representative confocal images show green fluorescence from anti-*hs*.circCAPN15*_bt_p* antibody staining, red fluorescence from Phalloidin staining of the cytoskeleton, and blue fluorescence from DAPI nuclear staining. Scale bar, 20 μm. **(C)** Western blot analysis of *CAPN15* protein (1,086 amino acids, ∼119.5 kDa) expression following *hs.*circCAPN15*_bt_p* knockout or overexpression (OE) in HeLa cells. *CAPN15* protein levels remained unchanged across all KO and OE conditions. The grayscale analysis of the bands was performed using ImageJ software. After normalization to β-tubulin, the relative protein level of *CAPN15* (*CAPN15*/tubulin) showed no statistically significant difference between the control and experimental groups (*p* >0.05, *n*=3). **(D)** Colony formation assay to evaluate the effect of *hs.*circCAPN15*_bt_p* KO and OE on HeLa cell proliferation. Each plate was seeded with 5,000 cells at the exponential growth phase, and cell density was measured after 14 days using a hemocytometer. **(E)** Wound-healing (scratch) assay to assess the impact of *hs.*circCAPN15*_bt_p* KO and OE on cell migration (*n*=3). Cells (5 × 10⁵ per well) were seeded in 6-well plates, scratches were introduced, and migration was monitored over 72 h. Scratch area was quantified using ImageJ. Migration rate was calculated as: (A - B)/A × 100%, where A is the initial scratch area (0 h), and B is the area at each subsequent time point. Statistical comparisons between groups were performed using Mann-Whitney U in GraphPad Prism (version 9.2.0). Statistical significance: *ns*: not significant, “*”: *p* <0.05, “**”: *p* <0.01, “***”: *p* <0.001, “****”: *p* <0.0001. Scale bar, 1,000 μm.

Subsequently, we adopted *Drosophila* to demonstrate how BT proteins function *in vivo*. Through systematic analysis of multi-source MS proteomes, we identified 248 putative circRNA-derived BT proteins (≥1 unique peptides per protein) potentially encoded by 1,251 sleep disorder-associated genes in *Drosophila*(*28*) (**Supplementary Table 7**). Among them, we selected five circRNAs for further experimental study: *dm.*circROLS*_bt_p*, *dm.*circFZ*_bt_p*, *dm.*circKUZ*_bt_p*, *dm.*circSK*_bt_p*, and *dm.*circTRPM*_bt_p*. RNase R-resistant RT-PCR confirmed circularity, and Sanger sequencing verified the BSJ (**fig. S7C and fig. S10A to D**). We employed the *elav*-GAL4 driver in the GAL4/UAS system(*29*) to achieve pan-neuronal overexpression of these five BT proteins from linear BW-ORF templates (**fig. S10E to I**), and subsequently assessed the rhythmic rest-activity patterns of *Drosophila* using established protocols(*30–32*). We found that overexpression of *dm.*circKUZ*_bt_p*, *dm.*circTRPM*_bt_p*, and *dm.*circFZ*_bt_p* in neurons did not significantly alter sleep patterns in either 3-day-old or 10-day-old *Drosophila* (**fig. S10J to L**), compared with the controls (*elav*-GAL4/+ and UAS-*dm.*circSK*_bt_p*/+, respectively). While overexpression of *dm*.circSK*_bt_p* increased the total nighttime sleep duration in 3-day-old *Drosophila*, with little effect on the number of nighttime sleep episodes and mean bout length, and enhanced the number of daytime sleep episodes in 10-day-old *Drosophila* without altering the total daytime sleep duration and mean bout length (**fig. S9M**). Sleep analysis of pan-neuronal *dm*.circROLS*_bt_p* overexpression revealed an increased number of daytime sleep episodes and decreased the daytime mean bout length in 3-day and 10-day *Drosophila* compared with the controls (*elav*-GAL4/+ and UAS-*dm*.circROLS*_bt_p*/+, respectively), indicative of sleep fragmentation, while total sleep duration remained relatively unaffected (**fig. S9N**). These preliminary observations implicated *dm*.circROLS*_bt_p* in sleep homeostasis, prompting further investigation. Of note, not all overexpressed BT proteins significantly altered sleep behavior in this initial screen, suggesting a degree of functional specificity for the phenotypes observed.

*ROLS* (rolling pebbles) is known to regulate myoblast fusion(*33*) and cell migration in *Drosophila*, while its human counterpart has been found implicated in intellectual developmental disorders (IDDs)(*34*). We then further characterized the two established *dm*.circROLS*_bt_p* KO lines (**Fig. 3H and I**). RT-qPCR showed no significant effects on endogenous linear *ROLS* or *dm*.circROLS transcription in the KO *Drosophila* (**Fig. 7A and B**). Immunofluorescence further revealed normal *ROLS* protein expression in larval salivary glands and adult brains in the KO lines (**Fig. 7C**), implicating the KO specificity of *dm*.circROLS*_bt_p*.

**Fig. 7.**
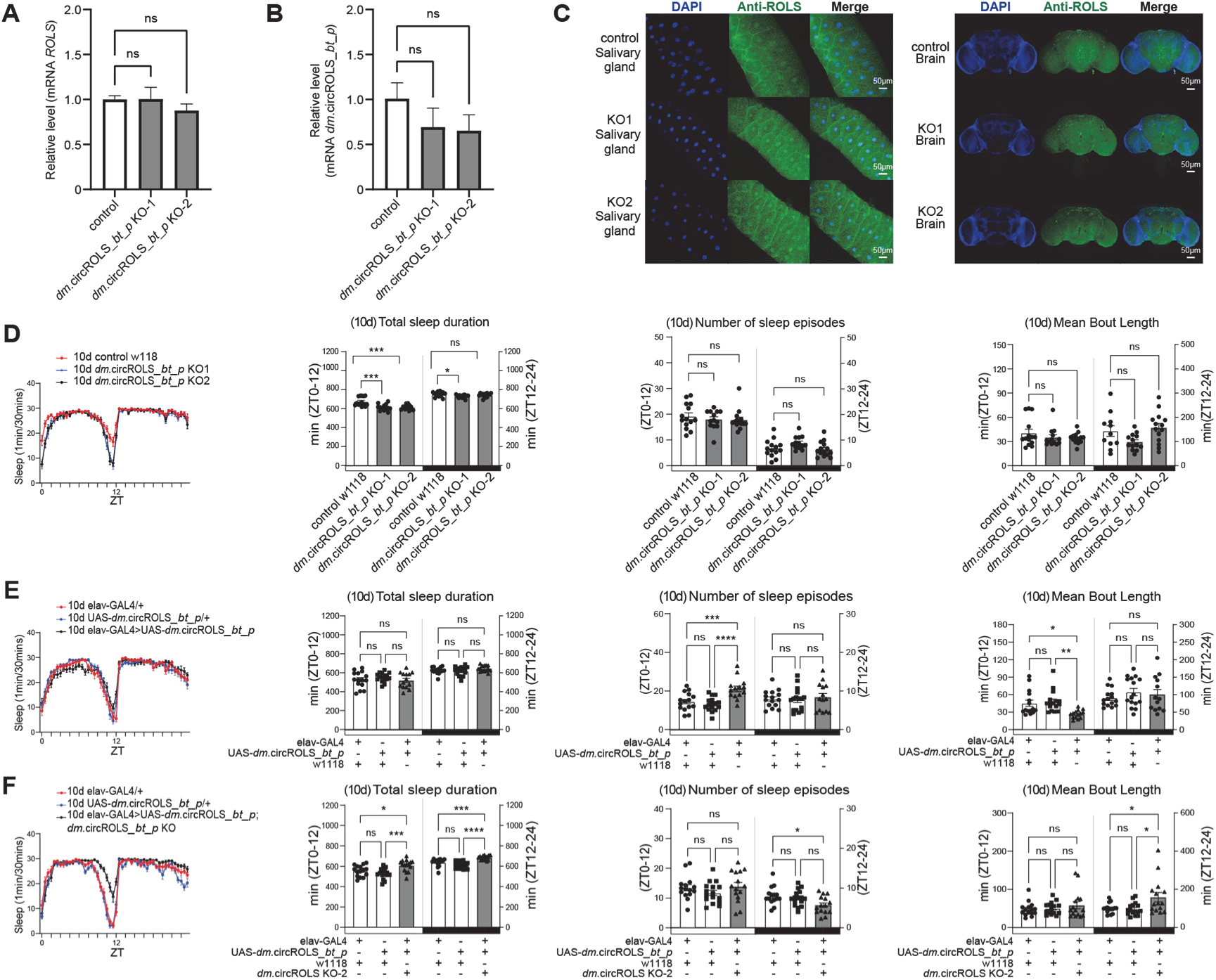
Overexpression of *dm*.circROLS*_bt_p* rescues the sleep phenotype caused by knockout. **(A)** Quantitative RT-qPCR analysis of parental *ROLS* mRNA expression in wild-type control and *dm.*circROLS*_bt_p* KO-1 and KO-2 *Drosophila* (*n*=3). **(B)** Quantitative RT-qPCR analysis of *dm*.circROLS expression in wild-type control, KO-1, and KO-2 *Drosophila*. All RNA samples were treated with RNase R to enrich for circular RNAs. Statistical comparisons between groups were performed using Mann-Whitney U (GraphPad Prism 9.2.0) with significance levels indicated as follows: *ns*: not significant, “*”: *p* <0.05, “**”: *p* <0.01, “***”: *p* <0.001, “****”: *p* <0.0001 (*n*=3). **(C)** Immunofluorescence analysis of parental *ROLS* protein expression in salivary glands of third instar larvae and adult brain of *Drosophila*, *dm*.circROLS *bt_p* KO-1 and KO-2, as well as w1118 (wild-type) control. Confocal microscopy images show *ROLS* protein detected with anti-ROLS antibody (green) and nuclei counterstained with DAPI (blue). Scale bar, 50 μm. **(D)** Sleep monitoring of 10-day-old *dm*.circROLS*_bt_p* KO-1, KO-2, and wild-type control. *Drosophila* were raised at 29 °C for 10 days prior to sleep analysis. **(E)** Sleep monitoring of 10-day-old *Drosophila* with pan-neuronal overexpression of *dm*.circROLS*_bt_p*. The experimental genotype was *elav*-GAL4/UAS-*dm*.circROLS*_bt_p*, and control genotypes were *elav*-GAL4/+ and UAS- *dm*.circROLS*_bt_p*/+. *Drosophila* were raised at 29 °C for 10 days before testing. **(F)** Sleep monitoring of 10-day-old *Drosophila* with pan-neuronal overexpression of *dm.*circROLS*_bt_p* in the *dm.*circROLS*_bt_p* KO-2 background. The experimental genotype was *elav*-GAL4/UAS*-dm.*circROLS*_bt_p; dm.*circROLS*_bt_p* KO/+. Control genotypes were *elav*-GAL4/+ and UAS-*dm.*circROLS*_bt_p*/+. **D-F**, from left to right, Sleep profiles depict 30-minute binned sleep accumulation over 24-hour periods. Total sleep duration represents the summed sleep time. The number of sleep episodes indicates a sleep bout. Mean Bout Length represents the mean sleep episode duration. An increase in the number of sleep episodes accompanied by a decrease in mean bout length indicates a sleep fragmentation phenotype. White and black bars along the x-axis indicate day (ZT0–12) and night (ZT12–24), respectively. Mann-Whitney U (GraphPad Prism 9.2.0) was used to statistically analyze the intergroup differences in mean bout length, with the following significance levels: *ns*: not significant, “*”: *p* <0.05, “**”: *p* <0.01, “***”: *p* <0.001, “****”: *p* <0.0001; (*n*>10). The other statistical comparisons between groups were performed using unpaired two-tailed t-tests (GraphPad Prism 9.2.0). Statistical significance: *ns*: not significant, “*”: *p* <0.05, “**”: *p* <0.01, “***”: *p* <0.001, “****”: *p* <0.0001; (*n*>10).

Sleep analysis showed that, exactly opposite to that of *Drosophila* overexpressing *dm*.circROLS*_bt_p* (**fig. S10N and fig. S11B**), *dm*.circROLS*_bt_p* KO 3-day-old adult displayed significantly reduced total daytime and nighttime sleep duration (vs. *elav*-GAL4/+, UAS-*dm.*circROLS*_bt_p*/+, *w*1118/+, *p* <0.01) (**fig. S11A**). Meanwhile, the sleep architecture (i.e., number of sleep episodes and mean bout length) was not significantly altered (**fig. S11A**). For 10-day-old adult *Drosophila*, the *dm.*circROLS*_bt_p* KO lines exhibited a consistent reduction in the sleep duration (vs. *elav*-GAL4/+, UAS-*dm.*circROLS*_bt_p*/+, *p* <0.01), without significant alteration of the number of sleep episodes and the mean bout length as well (**Fig. 7D**). Notably, pan-neuronal overexpression of *dm.*circROLS*_bt_p* in KO lines rescued the reduced total sleep duration in both 3-day-old (vs. *elav*-GAL4/+, UAS-*dm.*circROLS*_bt_p*/+, *p* <0.01) and 10-day-old (*p* <0.01) adult *Drosophila* (**Fig. 7F and fig. S11C**). Instead, both the total daytime and nighttime sleep duration increased when compared to the controls (*elav*-GAL4/+, UAS-*dm.*circROLS*_bt_p*/+) for 3-day and 10-day old adult *Drosophila*. The number of sleep episodes and mean bout length were less affected in this context (**Fig. 7E and fig. S11B**). These gene complement experiments further demonstrated that the observed sleep phenotype in the KO lines was indeed due to the elimination of *dm*.circROLS*_bt_p*, ruling out potential impacts of unintended destruction of host *ROLS* genes or non-specific effects during the knockout process. Taken together, these results concluded that *dm*.circROLS*_bt_p* is involved in *Drosophila* sleep regulation.

## Discussion

In eukaryotes, the canonical translation of linear mRNAs is primarily governed by 5’ cap structures and poly(A) tails(*35*). Under specific physiological stresses—such as those implicated in cancer, diabetes, and neurological disorders—cells may employ cap-independent initiation mechanisms mediated by internal ribosome entry sites (IRES) or other regulatory elements(*36*). Although these alternatives diversify translation initiation, they uniformly maintain 5’→3’ directionality, thereby reinforcing the long-standing dogma of unidirectional translation. This paradigm presupposes that all human proteins should be fully annotatable through mapping mass spectrometry (MS) data to the reference genome. However, a substantial fraction of acquired MS spectra remain unassigned and are often dismissed as background noise—a persistent discrepancy that points toward the existence of undiscovered coding mechanisms. Here, we provide evidence that eukaryotic circular RNAs can template 3’→5’ backward translation (BT), which may account for a portion of these uninterpreted spectra. Our integrated *in vivo* and *in vitro* analyses across human and *Drosophila* models establish BT as a biologically coherent process, distinct from annotation artifacts stemming from ribosomal scanning errors or computational noise. Notably, our analyses indicate that approximately 11.4% of circRNAs with coding potential can generate close to 1 million distinct protein products across species, as corroborated by MS or Ribo-seq data. This BT-encoded proteome substantially expands the known protein universe, revealing a previously cryptic layer of sequence diversity.

Backward translation produces polypeptides (BT proteins) that are typically concise yet bear distinct sequence and structural hallmarks, distinguishing them from canonical proteome products. Notably, a subset achieves cellular abundances rivaling those of annotated proteins, challenging their dismissal as stochastic translational noise. Genetic perturbations in human and *Drosophila* systems confirm this biological relevance: disrupting specific BT proteins yields discrete phenotypes without collateral effects on host gene expression or other circRNA functions (e.g., miRNA sponging(*37*)), underscoring their dedicated roles. Furthermore, BT appears amenable to regulation; bioinformatic signatures and engineered constructs suggest factors such as stem-loop motifs can influence BT expression. However, minimal synthetic motifs alone drive only marginal output, indicating that robust BT likely requires a synergistic interplay of *cis* elements, *trans* factors, and dynamic RNA architecture. Together—through unique molecular signatures, substantive expression, genetically encoded function, and regulatable capacity—BT is established as a coherent biological phenomenon rather than an analytical artifact. Its widespread occurrence across eukaryotes points to broad evolutionary relevance. Future efforts to decode its regulatory logic and functional proteome will further illuminate this hidden dimension of genetic information. Our findings thus elevate BT to a bona fide pathway for proteomic innovation, revealing a long-overlooked source of functional diversity within eukaryotic cells.

A central question posed by our discovery is how the translation machinery operates in the 3’→5’ direction on circRNA templates. Our data suggest that backward translation may, in part, adhere to canonical principles—utilizing triplet codons and requiring a defined start codon. Furthermore, we find that IRES elements and initiation sites from BT-active circRNAs can support both forward and backward translation, implying shared initiation factors. While the circular template likely imposes unique topological constraints that facilitate bidirectional movement, the notion of 3’→5’ elongation remains mechanistically challenging. Precedents such as (–1) ribosomal frameshifting(*38*) and factor-mediated back-translocation in prokaryotes and organelles(*39*)^,^ (*40*), however, indicate that limited reverse movement is possible within the translational apparatus. A plausible mechanistic model involves a structural reorientation of the ribosome, enabling it to engage the circRNA in a symmetrically flipped manner (**Fig. 1A**). This view is supported by established precedents for reverse Watson–Crick (rWC) base pairing, which provides a structural basis for parallel-strand nucleic acid interactions(*^41, 42^*). In an rWC geometry with trans glycosidic bonds, codon– anticodon pairing may occur on a circRNA template (**fig. S1A**), allowing for inverse substrate recognition without necessitating a complete reassembly of the core ribosome. Taken together, while the detailed mechanism of BT awaits full resolution, these functional and structural insights suggest that the translational machinery possesses a latent capacity for directionality that extends beyond the canonical 5’→3’ paradigm.

There are two relevant issues worth mentioning regarding the results. First, our current research focuses on 5’-GUA-3’ initiating BT; however, we cannot rule out the possibility of the occurrence of non-5’-GUA-3’ initiating BT(*43*). Second, this study suggests that circRNAs are the primary template for BT, as to date, we do not have any clear evidence that linear mRNAs are capable of initiating BT despite efforts that have been made (data not shown). We also cannot rule out the possibility that linear mRNAs or circRNAs may undergo sequence reversal through multiple rounds of transcription or reverse transcription, thereby producing protein products resembling those from backward translation. However, to date, such backward transcription phenomena and key enzymatic machinery required to catalyze such a process have not been reported.

In summary, our findings provide evidence for a backward translation mechanism mediated by circRNAs, which appears to expand the coding potential of genomes and uncover a layer of proteomic diversity. The functional significance and distinctive features of BT-generated proteins merit future investigation, as they may hold broad biological and therapeutic implications.

## Acknowledgement

We are extremely grateful to the distinguished scholars across multiple biological disciplines—including Prof. Mu-Ming Poo, Prof. Senfang Sui, Prof. Juergen Brosius (despite his continued skepticism), Prof. Qinxi Li, Prof. Aidong Han, Prof. Bingwei Lu, Prof. Xinping Yang, Prof. Li-Zhi Mi, Prof. Guixuan Zhou—for their insightful guidance and valuable assistance in the course of preparing this paper. We thank Dr. Xuan Guo and Dr. Wei Wu for *Drosophila* transgenic and gene editing assistance, Tsinghua *Drosophila* stock center for stocks. We thank Professor Xian Zen and Dr. Hui Zhao for assistance in circRNA synthesis. We thank Yaying Wu, Zheni Xu and Dr. Changchuan Xie for mass spectrometry experiments and data analysis.

## Contributions

Conceptualization: Zhi-Liang Ji, Zhe Lin Methodology: Zhi-Liang Ji, Yufeng Yang, Zhe Lin Formal analysis: Ziwei Lv, Zhe Lin, Huajun Yin, Muxi Li, Juan Du, Shiyu Wang Investigation: Xiangyou Pi, Dani Shi, Tianliang Liu, Yanchao Wang, Peng Wang, Wenfeng Chen, Ling Sun, Wenbo Lin, Tao Tao Software: Zhe Lin, Yangmei Qin Writing: Zhi-Liang Ji, Yufeng Yang, Zhe Lin, Ziwei Lv, Xiangyou Pi Project administration: Zhi-Liang Ji, Yufeng Yang

## Data and materials availability

The circRNA sequences, the BT protein sequences, and the self-coded scripts used in this study are all freely accessible at GitHub (https://github.com/BADDxmu/BawaCirc).

## Supplementary Materials Materials and Methods

### Data collection and preprocessing

In all, 379 transcriptomes were obtained from the National Center for Biotechnology Information (NCBI) SRA, spanning six species. The raw Tandem Mass Spectrometry (MS/MS) files of 6,112 proteomes were obtained from the ProteomeXchange (ProteomeCentral Datasets)(*44*) and the PRIDE (Proteomics Identification Database)(*45*). Besides, 493 Ribosome Profiling (Ribo-seq) translatomes were obtained from the NCBI SRA. These datasets were cataloged in **Supplementary Table 1**. All transcriptome datasets were processed for adapter trimming and contaminant removal using Trimmomatic (version 0.38)(*46*), with adapter sequences provided either by the data source or the corresponding sequencing platform. Quality assessment of the trimmed reads was performed using FastQC (https://www.bioinformatics.babraham.ac.uk/projects/fastqc/).

### CircRNA detection

The circRNAs were detected from the RNA-seq transcriptomes with the CIRIT software(*19*) and the full-length sequences of circRNAs were generated as well. In addition, a collection of 1,532,085 circRNAs and their sequences was sourced from several prominent circRNA databases, including Circbank(*47*), CircAtlas(*48*), CircBase(*49*), and TransCirc(*16*). All circRNAs were annotated with the self-developed trCIRIT-BSJ module via blasting the reconstructed full-length sequences to the reference genome using the gmap tool (version 2020-10-14)(*50*), whereby the exon-intron structure of circRNAs as well as the back-splicing junction (BSJ) sites were determined. Unless otherwise specified, all genomes were as the following versions: *Saccharomyces cerevisiae* (R64), *Caenorhabditis elegans* (WBcel235), *Drosophila melanogaster* (Release 6 plus ISO1 MT), *Danio rerio* (GRCz11), *Mus musculus* (GRCm39), and *Homo sapiens* (T2T-CHM13). The expression levels of circRNAs were quantified using the RSEM(*51*) tool. For quantification, a self-coded script was used to extract 140 nucleotides (nt) upstream and downstream of each circRNA BSJ site, which was then aligned with the raw sequence data. The detailed step-by-step operating procedure is publicly available at GitHub (https://github.com/BADDxmu/BawaCirc).

### Detection of circRNA translation with mass spectrometry

The methodology of circRNA translation detection from mass spectrometry (MS) data was schematically illustrated in **fig. S2 and fig. S4**. The circRNA sequences were first re-coordinated according to the circRNA annotation and unfolded to linear sequences at the BSJ sites. The linear sequences were then expanded three times and joined together in a head-to-tail way. The expanded sequences were fed into the trCIRIT-ORF module for open reading frame (ORF) prediction. The trCIRIT-ORF module employs the regular expression “(?=(ATG(?:(?!TAA|TAG|TGA)…)*(?:TAA|TAG|TGA)))” to identify all open reading frames (ORFs) within circRNA sequences that begin with an ATG start codon and terminate with a TAA, TAG, or TGA stop codon. For the backward ORFs, the sequences were reversed before input for ORF prediction.

The predicted ORFs served as the reference database for peptide search from the MS/MS spectrometry files with the MaxQuant software v.2.0.2.0(*52*). Before peptide search, the BW ORFs were aligned against the corresponding genomes to eliminate putative palindromic sequences (sequence similarity >80%) with Blastn. For peptide search with MaxQuant, trypsin was selected as the proteolytic enzyme, and two missed cleavage sites were allowed. The first search mass tolerance was set to 20 ppm, and the main search peptide tolerance was set to 4.5 ppm. The false discovery rates (FDR) for peptide-spectrum matches (PSMs) and proteins were set to less than 1%. The iBAQ data from MaxQuant were normalized according to the method described in Jiang, Y. *et al*(*23*). The detected unique peptides were further aligned against UniProtKB to exclude those shared peptides that originated from linear mRNAs (sequence similarity = 100%). The remaining unique peptides then served as mass spectrometric proof of circRNA translation. The detailed step-by-step operating procedure is provided in **Supplementary Text** and is also publicly available at GitHub (https://github.com/BADDxmu/BawaCirc).

### Detection of active circRNA translation with ribosome profiling analysis

The methodology of circRNA translation detection from ribosome profiling/sequencing (Ribo-seq) data were schematically illustrated in **fig. S3**. The active backward translation events were detected by aligning the preprocessed Ribo-seq reads exclusively to the BSJ and its flanking 25nt sequences (A total of 50nt) spanned by the backward ORFs with Bowtie v1.0.0(*53*) (parameter -v 1), excluding those for the rRNA, linear mRNA transcripts (GRCh38.p14, Feb 3, 2022), ncRNAs (the RNAcentral database(*54*)) and circRNA forward ORFs one by one (parameter -v 3). The retained reads were converted to the BAM files with samtools v.1.3.1(*55*), which were subsequently interpreted and analyzed with bamdst v.1.0.9 (https://github.com/shiquan/bamdst) and mosdepth v.0.3.10 (https://github.com/brentp/mosdepth). The BAM files were also used for later investigation of backward translation characteristics. The detailed step-by-step operating procedure is provided in **Supplementary Text** and is also publicly available at GitHub (https://github.com/BADDxmu/BawaCirc).

### Diagnostic analysis of ribosome profiling

The quality of Ribo-seq data was further diagnosed using RiboWaltz(*56*), which helps determine P-site offsets, evaluate 3-nt periodicity, and visualize metagene read distributions. A publicly available dataset (SRR6838651) provided by the RiboWaltz developers was used as a reference dataset for conventional translation of linear mRNAs. For backward translation analysis, we first used Bowtie v1.0.0 (parameter: -v 3) to remove Ribo-seq reads that could be aligned to linear mRNAs, ncRNAs, and forward ORFs of circRNAs. The remaining unmapped reads were aligned to backward ORFs of circRNA (Bowtie v1.0.0, parameter: -v 3) and subsequently subjected to RiboWaltz analysis. The detailed step-by-step operating procedure is publicly available at GitHub (https://github.com/BADDxmu/BawaCirc).

### Statistics

Linear regression models in **fig. S6B and C** were fitted using the lm function from the stats package (version 4.3.3) to compute the coefficient of determination (*R*^2^), a measure of how well the regression line explains the variability in the dependent variable.

The *R*^2^ value is calculated as:

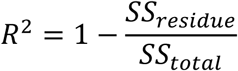

Where *SS*_residue_ is the sum of squares of residuals, and *SS*_total_ is the total sum of squares. An *R*^2^ value of 1 indicates a perfect fit, meaning all data points lie on the regression line, while a value of 0 means the model does not explain any variability. Values closer to 0 suggest a weaker relationship between the variables. The sample size for the analysis is reported in the corresponding figure legends.

Unless otherwise stated, intergroup comparisons were performed using unpaired two-tailed t-tests (for *n* ≥ 10) or Mann-Whitney U tests (for *n* < 10), implemented in GraphPad Prism 9.2.0 with significance levels indicated as follows: “*” *p <*0.05; “**” *p* <0.01; “***” *p* <0.001; “****” *p* <0.0001. “*ns*” indicates no significant difference (*p* >0.05). Damaged samples were excluded from immune imaging analyses.

### Translation initiator sequence analysis

The potential IRES-like sequences in circRNA were detected with IRESfinder software v.1.1.0(*26*). The IRES-like 6nt at the 20nt flanking sequences preceding the backward start codon (GUA) were detected by Blastn (v.2.5.0), allowing no mismatch. The Kozak sequence (reversed: GGUACCACCA) was checked with a self-coded script. The m^6^A modifications were predicted with the local tool of the SRAMP website(*57*).

### Domain and motif analysis of BT proteins

The domain information of BT proteins was obtained via a search of the SMART database(*21*), covering the SwissProt, the SP-TrEMBL, and the stable Ensembl protein sets. The motif analysis was demonstrated with the tool ps_scan (version 2018-01-25), referring to the PROSITE database(*22*) (prosite.dat, version 2024-03-27). A custom script was employed to calculate amino acid usage frequencies and dipeptide frequencies. The script is publicly accessible at GitHub (https://github.com/BADDxmu/BawaCirc).

### RNA Secondary Structure Prediction

The RNAfold web server (http://rna.tbi.univie.ac.at/cgi-bin/RNAWebSuite/RNAfold.cgi) was utilized to predict the secondary structure of circRNAs with the option “assume RNA molecule to be circular”. The secondary structure with MFE (Minimum Free Energy) was chosen for presentation.

### Protein structure simulation and clustering

The structures of selected BT proteins were modelled with the AlphaFold2 web service(*58*). To obtain a stable conformation, a 50ns molecular dynamics (MD) simulation was conducted with the Gromacs software(*59*), adopting the GROMOS 54 force field and the TIP3 water environment. The final frame of the MD simulation was taken as the protein optimized structure. The protein structure was visualized with ChimeraX (v1.7).

For structure clustering, a total of 542,337 known protein sequences and corresponding structures were downloaded from the AlphaFold website (https://alphafold.com/download). Subsequently, CD-HIT v4.7 was utilized to merge the similar protein sequences (parameters: -c 0.4 -n 2) to reduce the computational cost. Before clustering, the structures of both the known proteins and the selected BT proteins were represented as the average embedding tokens with the ProtT5-XL-U50 model of ProtTrans(*60*). The K-means method was utilized to cluster the structures. The t-SNE method of the cuML package (v22.06.01) was utilized to visualize the clustering result. Matplotlib (v3.5.3) and Seaborn (v0.11.2) are used to visualize the result in reduced dimensions.

### Mass Spectrometry

*Drosophila melanogaster* (w1118) were homogenized on ice in pre-chilled RIPA lysis buffer (Beyotime, P0013B) using an electric homogenizer. Homogenates were incubated on ice for 30 min and then centrifuged at 12,000 rpm for 10 min at 4°C. The resulting supernatants were collected, and total protein concentrations were determined using a BCA protein assay kit (Beyotime, P0010).

For immunoprecipitation (IP), Protein A/G magnetic beads (Beyotime, P2018) were equilibrated by washing 4–6 times with 1× TBS buffer (50 mM Tris-HCl, 150 mM NaCl, pH 7.4). The beads were then incubated with 5–10 μg of anti-*dm.*circROLS*_bt_p* antibody (peptide epitope: DSNIKQKFKRKINTC) at room temperature for 1 h with gentle agitation. After antibody coupling, the beads were washed 4–6 times with TBS to remove unbound antibodies. Subsequently, 200–500 μg of protein lysate was added to the antibody-conjugated beads and incubated overnight at 4°C with gentle rotation. The next day, beads were washed 5–6 times with TBS to eliminate nonspecifically bound proteins.

Elution of bound proteins was performed with freshly prepared 8 M urea in 0.1 M Tris-HCl (pH 8.0) at room temperature for 5 min. The eluted proteins were collected, and concentrations were re-quantified using the BCA assay. Proteins were subjected to in-solution tryptic digestion, followed by drying under vacuum.

Peptides were resuspended in 10 μl of 0.1% formic acid and analyzed using an EASY-nLC 1200 system (Thermo Fisher Scientific) coupled to an Orbitrap Fusion Lumos mass spectrometer (Thermo Fisher Scientific) equipped with an EASY-IC ion source. Samples were loaded onto a homemade C18 column (35 cm × 75 μm i.d., 2.5 μm, 100 Å) and eluted over 120 min using a linear gradient of 3–35% acetonitrile in 0.1% formic acid at a flow rate of 300 nl/min.

For data-independent acquisition (DIA), a 25 m/z isolation window spanning 400– 1200 m/z was used. DIA data were processed using DIA-NN (v1.8.1)(*61*), searched against a custom *Drosophila* UniProtKB protein database supplemented with the *dm.*circROLS*_bt_p* sequence. The analysis was performed in library-free mode with deep learning-based spectral and retention time prediction enabled. Search parameters included: protease, Trypsin/P; maximum of two missed cleavages; N-terminal methionine excision, enabled; fixed modification, carbamidomethylation (C); variable modification, oxidation (M). Mass accuracy tolerances were set to 10 ppm for both MS1 and MS2. Protein inference was based on protein names from the FASTA file, and quantification was conducted in “Robust LC (high accuracy)” mode. All other settings were kept at the default.

Raw MS data have been deposited and are publicly accessible at GitHub: https://github.com/BADDxmu/BawaCirc.

### Cell culture

HEK-293T and HeLa cell lines were authenticated by STR analysis. Cells were cultured in DMEM medium (Gibco, C11995500BT), supplemented with 10% fetal bovine serum (Gibco, ST30-3302) and 1% penicillin-streptomycin (Beyotime, 15140-122), and incubated at 37°C in a humidified incubator with 5% CO_2_.

### RT-PCR/RT-qPCR and Sanger Sequencing

Total RNA was extracted from *Drosophila* (w1118) and HeLa cells using RNAiso Plus (TaKaRa, 9108) according to the manufacturer’s instructions. The total RNA was homogenized thoroughly, and the gDNA was removed using DNase I. RNase R (Beyotime, R7092S) treatment was performed to degrade linear RNA following the manufacturer’s instructions. The cleaned circRNA was reverse-transcribed into cDNA using a reverse transcription kit with random hexamers (Thermo Scientific, K1622). Divergent primers were designed (**Supplementary Table 4**) to amplify the sequences spanning the circRNA BSJs with PCR. The amplified products were analyzed by electrophoresis on a 1% agarose gel. Bands of the expected size were excised and purified using the SanPrep Column DNA Gel Extraction Kit (Sangon Biotech, B518131). Sanger sequencing (Tsingke Biotech, Fujian) was utilized to determine the sequence composition of purified DNA. Meanwhile, the RT-qPCR primers were designed to quantify circRNAs (**Supplementary Table 4**).

### Immunofluorescence assay

The antigenic epitopes were designed based on the backward ORF regions of circRNAs (**Supplementary Table 8**). These short epitope peptide fragments were synthesized and injected into rabbits to generate antibodies, which were subsequently isolated and purified (GenScript, Nanjing). The following reagents were also utilized for different purposes in this study: Alexa FluorTM 488 goat anti-rabbit IgG (H+L) (Invitrogen, A11008), Alexa FluorTM 568 goat anti-mouse IgG (H+L) (Invitrogen, A11031), and GF®568-Phalloidin (GenXion, JXF40131).

For immunofluorescence, cells were seeded in a 24-well covered plate and cultured for 24 hours in an incubator with 5% CO_2_. Cells were then fixed with 4% paraformaldehyde for 10 minutes and washed 2-3 times with PBS. Permeabilization was performed with 0.2% Triton-X 100 for 1 minute, followed by 2-3 PBS washes. Cells were incubated with 2% bovine serum albumin (BSA) at room temperature for 1 hour, and then with primary antibodies overnight at 4°C. After primary antibody incubation, cells were washed 2-3 times with PBS, 5 minutes each wash, and then incubated with secondary antibodies at room temperature for an hour. Following secondary antibody incubation, cells were washed 2-3 times with PBS and incubated with DAPI and/or phalloidin for 10 minutes each. After 2-3 PBS washes, an anti-fade reagent was added to the 24-well plate, and coverslips were mounted onto slides using tweezers. Imaging was performed using a confocal microscope.

For immunofluorescence assay in *Drosophila*, the third instar larvae were fixed in 4% PFA at room temperature for 2 hours. After fixation, the larvae were washed three times with PBST (PBS + 0.4% Triton X-100) and then dissected in PBST to isolate the brain, eye discs, wing discs, and salivary glands. The samples were blocked at room temperature for 1.5 hours in a blocking solution consisting of 950 μL PBST and 50 μL normal goat serum. After removing the blocking solution, the samples were incubated with primary antibody solution (950 μL PBST, 50 μL normal goat serum, and 2 μL primary antibody) at 4°C for 24 hours. Following primary antibody incubation, the samples were washed three times with PBST and then incubated with secondary antibody (Invitrogen: Alexa Fluor™ 488 goat anti-rabbit IgG (H+L)) at 4°C for 48 hours. After recovering the secondary antibody and washing three times with PBST, the samples were incubated with DAPI (Beyotime, C1002) at room temperature for 20 minutes. Finally, the samples were washed three times with PBST and mounted using anti-fade mounting medium (Beyotime, P0126). For detecting apoptosis signaling in *Drosophila*, the Cleaved Caspase-3 (D175) Rabbit antibody (Cell Signaling, 9661S) was used.

### Western blotting

Samples were collected on dry ice. Heads were homogenized in RIPA lysis buffer (Beyotime, P0013B) using a motorized pestle. Lysates were then centrifuged at max speed for 10 min, and the supernatant was saved (12,000 rpm). All lysates were boiled with protein sample buffer. Proteins were quantified using the BCA Protein Assay Kit (Beyotime, P0010), and 20 mg of protein was loaded on pre-stained SDS-PAGE gels (Bio-Rad). Proteins were transferred to 0.45 mm Nitrocellulose Membranes using wet transfer for 60 min. Unspecific binding was blocked using 5%BSA powder in TBST (BioFroxx, 4240GR100). Primary antibodies were diluted and incubated with the membrane overnight at 4℃. HRP-coupled secondary antibodies were used according to the primary antibody. Signal was developed using ECL Western Blotting Detection Reagents (Beyotime, P0018S). β-Tubulin was used as a normalization control. The following reagents were also utilized for different purposes in this study: β-Tubulin Mouse Monoclonal Antibody (Beyotime, AF2835), HRP-labeled Goat Anti-Mouse IgG(H+L) (Beyotime, A0216), and HRP-labeled Goat Anti-Rabbit IgG(H+L) (Beyotime, A0208).

### Plasmid cloning and transfection into human cells

The plasmids for human cell lines in this study were generated using pcDNA3.1(+) ZKSCAN1 MCS + Sense IRES (Addgene: #69909) and pcDNA3.1(+). The plasmid includes a CMV and a T7 promoter that drive the transcription of an upstream intron, the expected ORF, and a downstream intron. The introns form a circular RNA through complementary base pairing, which subsequently expresses the target ORF. The compositions of plasmids for human cell transfection designed in this study are illustrated in **Fig. 4A**. The plasmids were transfected into HEK-293T or HeLa cells via an electroporation kit (Celetrix, 1216) according to the experimental needs. For electroporation of HEK-293T cells, the voltage was set to 600V with a duration of 30ms. Six hours post-electroporation, the medium was changed to remove the electroporation solution. The backward translation initiation sites (TISs) used in this study are compiled in **Supplementary Table 8**.

### Circular RNA synthesis and validation

Scarless circRNAs were synthesized via a permuted intron‒exon system (PIE)-based circularization strategy with the assistance of Byterna Therapeutics Ltd (Shanghai, China). Synthetic sequences listed in **Supplementary Table 9**, including type I intron fragments, were cloned into the pUC57 plasmid containing a T7 promoter. The resulting plasmids were linearized using restriction endonucleases and subjected to in vitro transcription with the T7 RNA polymerase system (New England Biolabs, E2040L). Residual DNA was removed using DNase I (New England Biolabs, M0303L). Following transcription, RNAs were purified using phenol: chloroform: isoamyl alcohol (25:24:1) extraction to eliminate proteins and enzymatic contaminants.

To enhance RNA stability and cyclization efficiency, polyadenylation was performed using *E. coli* Poly(A) Polymerase (New England Biolabs, M0276M) in the presence of ATP at 37 °C, typically resulting in the addition of 100–200 adenosine residues to the 3’ end of the RNA. Polyadenylated transcripts were further purified via ethanol precipitation.

Purified RNAs were circularized in a 20 μL reaction containing 1× T4 RNA ligase buffer (New England Biolabs, B0216L) and 2 mM GTP at 55 °C for 15 minutes. The reaction was terminated with 2 μL of 100 mM EDTA (pH 8.0), and 2 μL of 100 mM MgCl₂ was added to neutralize the EDTA and restore downstream enzymatic activity. The resulting circRNAs were treated with RNase R (Novoprotein, E224) to degrade linear RNAs and subsequently purified using the Monarch Spin RNA Cleanup Kit (New England Biolabs, T2040S).

### Validation of circularization

To confirm RNA circularization at the splice junction, RNase R-treated RNA was reverse transcribed using the RevertAid First Strand cDNA Synthesis Kit with random primers (Thermo Fisher Scientific, K1622). PCR was then performed using primers spanning the back-splice junction, and the amplicons were validated by Sanger sequencing.

### Immunoprecipitation of GFP protein

HEK-293T cells expressing GFP fusion proteins and non-transfected controls were harvested by centrifugation (1300 rpm, 5 min, 4 °C) and washed twice with ice-cold PBS. Cells were lysed on ice for 30 minutes in NP-40 lysis buffer (50 mM Tris-HCl, pH 7.4, 150 mM NaCl, 1% NP-40, 10% glycerol), and lysates were cleared by centrifugation (12,000 rpm, 10 min, 4 °C). The supernatant was collected, and protein concentrations were determined using a BCA assay.

Protein A/G magnetic beads (Beyotime, P2018) were washed 4–6 times with 1× TBS and incubated with 5–10 μg of anti-GFP antibody (Thermo Fisher, A-11122) or isotype-matched IgG control for 1 hour at room temperature. After washing, 200– 500 μg of protein lysate was added to the antibody-coupled beads and incubated overnight at 4 °C on a rotator. The beads were washed 5–6 times with 1× TBS and resuspended in 50 μL of non-reducing 1× SDS-PAGE sample buffer. Samples were boiled for 10 minutes, and 25 μL of the supernatant was analyzed by Western blotting.

### In vitro translation assay

To assess backward translation from circRNAs, we used the Rabbit Reticulocyte Lysate System (Promega, L4960) for Gaussia protein synthesis mediated by the TIS element. In vitro translation was performed with 1.5 μg of RNA incubated at 30 °C for 120 minutes according to the manufacturer’s instructions. Luminescence was measured using the Gaussia-Lumi™ Luciferase Reporter Assay Kit (Beyotime, RG072S). For detection, Gaussia-Lumi™ substrate and buffer were mixed at a 1:100 ratio to prepare the working solution. A total of 10 μL of translation product was mixed with 100 μL of detection reagent (final volume: 110 μL), incubated for 5 minutes at room temperature in the dark, and chemiluminescence was recorded using the GloMax Multi Jr detection system (Promega, E6070) with an integration time of 10 seconds per well.

### *Drosophila* Translation System

#### Preparation of *Drosophila* Ovarian Extract

The *Drosophila* ovarian extract was prepared according to previously described methods with minor modifications(*62*). Briefly, young female flies were cultured at 25°C on a yeast-containing diet for 2 to 3 days. Ovaries were dissected in ice-cold PBS and collected. All subsequent steps were performed on ice or at 4°C. The collected ovaries were allowed to settle by gravity, and the packed volume was measured. The ovarian pellets were then washed twice with a 12-volume mixture of PBS and Extraction Buffer DEI (10 mM Hepes pH 7.4, 5 mM DTT, supplemented with 1X COMPLETE-Protease Inhibitors Cocktail EDTA-free from Roche) at a 1:1 ratio. This was followed by two quick washes with a 12-volume of DEI buffer alone. The washed ovaries were transferred to a Dounce homogenizer (Kimble, 885300-0002). After removal of excess buffer, the ovaries were directly homogenized using the tight pestle. The homogenate was transferred to a microcentrifuge tube and centrifuged at 14,000 rpm for 10 minutes at 4°C. The resulting supernatant was carefully collected, representing the ovarian extract ready for subsequent in vitro translation assays. The extract could be aliquoted and stored at - 80°C or used immediately.

#### In Vitro Translation

The in vitro translation was performed based on literature protocols with conditions optimized for circular RNA(*62, 63*). In a 25 µL reaction mixture, 1 µg of template circular RNA was translated in a system containing 60 µM amino acid mixture, 16.8 mM phosphocreatine, 80 ng/mL creatine kinase, 24 mM HEPES pH 7.4, 0.6 mM magnesium acetate, 60 mM potassium acetate, 0.1 mM spermidine, 1.2 mM DTT, 100 µg/mL bovine liver tRNA, and 40% ovarian extract. The reaction was incubated at 25°C for 120 minutes. Luciferase production was detected using a chemiluminescence assay (Beyotime, RG072S). For detection, Gaussia-Lumi™ substrate and buffer were mixed at a 1:100 ratio to prepare the working solution. A total of 10 μL of translation product was mixed with 100 μL of detection reagent (final volume: 110 μL), incubated for 5 minutes at room temperature in the dark, and chemiluminescence was recorded using the GloMax Multi Jr detection system (Promega, E6070) with an integration time of 10 seconds per well.

### Knock-in of red fluorescent protein

To knock-in a fluorescent protein into circRNA, we inserted a reversed linker-mScarlet3 sequence just upstream of the backward stop codon (GAT) (**Fig. 3A**). For *hs*.circCAPN15, we identified the target site within 30 bp of the backward stop codon (GAT) of *hs*.circCAPN15 and designed sgRNAs targeting 5’-CCACCGGATAGTTA-3’ and 5’-CACCCGGATAGTTT-3’. These sgRNA sequences were cloned into pSpCas9 (BB) -2A GFP (PX458) (Addgene: # 48138), digested with the BbsI enzyme. The donor plasmid consists of homologous arms (800-1000 bp) located on the GAT site flanks, with reversed linker (5’-GGAGGTGGAGGTAGTGGTGGAGGAGGTAGTGGTGGTGGAGGTTCT -3’) and reversed mScarlet3 sequences inserted in appropriate directions between the homologous arms, and cloned into the pBlueScript II SK (+) plasmid. Extract plasmids and transfect them into HeLa cells through electroporation. 48 hours later, flow cytometry was performed on the cells using red fluorescent protein mScat3HO (Becton Dickinson, FACSARia III). Single cells were sorted into 96-well plates, and amplified monoclonal cells were identified by sequencing.

For *dm*.circROLS, we identified the target site within 30 bp of the backward stop codon (GAT) of *dm*.circROLS and designed sgRNAs targeting 5’-CCAGTTTGGCCCCCTTGCGC-3’ and 5’-ACAGGTGTTCCTGCGCAAAG-3’.

These sgRNA sequences were cloned into pU6-BbsI-chiRNA (Addgene:#45946). The donor plasmid consists of homologous arms (800-1000bp) located on the TAG site flanks, with reverse linker and reverse mScarlet3 sequences inserted in appropriate directions between the homologous arms, and cloned into the pBlueScript II SK (+s) plasmid. After the construction of the above plasmid, it was injected into the *Drosophila* egg by microinjection (BL78781).

### Construction of CRISPR knockout cell lines

Target gene sites for *hs.*circCAPN15 were identified using CCTop - CRISPR/Cas9 target online predictor, resulting in two target sequences: 5’-GTAGGCAGGGGAGGTAGCAG-3’ and 5’-CTACCTCCCCTGCCTACCTT-3’.

These target sequences were then cloned into the pSpCas9(BB)-2A-GFP (PX458) vector (Addgene: #48138), which had been digested with BbsI. The plasmids were extracted and transfected into HeLa cells via electroporation. After 48 hours, GFP-positive cells were sorted using fluorescence-activated cell sorting (FACS) with a FACSAria III sorter (Becton Dickinson, FACSAria III). Single cells were collected into 96-well plates, and the resulting clonal cell lines were expanded and sequenced for validation.

### Generation of stable *hs*.circCAPN15 overexpression cell lines

To express the *hs*.circCAPN15*_bt_p* protein via conventional forward translation, we synthesized the *hs*.circCAPN15 backward ORF in the forward orientation. The synthesized fragment was digested with NheI and BamHI, then ligated into a proprietary Celetrix selection vector (unpublished). Plasmid constructs were prepared using the ZymoPURE II Plasmid Midiprep Kit (ZYMO RESEARCH, D4201).

HeLa cells in optimal growth condition were harvested, re-suspended in Celetrix electroporation buffer (Celetrix, 1216), and electroporated with the *CAPN15* selection plasmid (640V, 30ms). Post-electroporation, cells were plated in 6-well plates, with 3 ml of complete medium added per well. After three days of incubation at 37°C, cells were sub-cultured when they reached approximately 80% confluence. Selection was performed using puromycin at concentrations of 1.0 µg/ml, 2.0 µg/ml, and 5.0 µg/ml. Cells that survived two weeks of selection under 2.0 µg/ml puromycin were expanded, and successful overexpression was confirmed by qPCR and immunoblotting.

### Cell scratch

Cells were digested using 0.25% trypsin, and an appropriate amount was transferred to a hemocytometer for cell counting. Cells were seeded into a 6-well plate at a density of 5×10^5^ cells per well. Three parallel lines with equal spacing were drawn on the back of each well using a marker. Once the cells reached 80-90% confluence, a 200 μL pipette tip was used to draw three vertical lines perpendicular to the parallel lines on the bottom of the plate, using a ruler for guidance. Cell migration was observed under an inverted microscope every 12 hours. The scratch area at each time point was quantified using ImageJ software, and the cell migration rate was calculated using the formula: (A−B)/A×100%, where A is the scratch area at 0 hours and B is the scratch area at subsequent time points.

### Plate colony formation assay

Exponentially growing cells were harvested and digested into single cells using 0.25% trypsin. After cell counting with a hemocytometer, 5000 cells were seeded into each well of a six-well plate and a 10 cm culture dish. The cells were gently shaken to ensure even distribution and then incubated at 37°C with 5% CO_2_. Colony formation was assessed on days 2 and 7 for the six-well plates, and on day 14 for the 10 cm dishes. At the end of the incubation period, the medium was aspirated, and the cells were washed twice with PBS. The cells were then fixed with 4% paraformaldehyde for 15 minutes, washed twice with PBS, and stained with crystal violet for 15 minutes. Excess stain was removed by washing with running water three times. The plates were air-dried and photographed. The number of colonies formed from 100 single cells was counted and averaged to determine the colony formation efficiency.

### Construction of a CRISPR knockout *Drosophila* strain

*dm*.circROLS is derived from exons 5, 6, and the 3’UTR of its host gene *ROLS* through back-splicing. This circular RNA contains an open reading frame (ORF) encoding a 114 amino acid peptide, with its reverse initiation codon (GTA) overlapping the host gene’s 3’UTR region.

Using the CCTop CRISPR/Cas9 target online predictor, we identified target sites within 30 bp of the *dm*.circROLS backward start codon (GTA) and designed sgRNAs targeting 5’-GGAATAGCGATTAATTCGAA -3’ and 5’-ATAGCAAAAATTAAGGTCTA -3’. These sgRNA sequences were cloned into the BbsI-digested pU6-BbsI-chiRNA vector (Addgene: #45946).

To achieve the desired mutation, we changed the GTA backward start codon of *dm*.circROLS to GTC. The donor plasmid was constructed by cloning 800-1000 bp homology arms flanking the mutated GTA site into the KpnI and ScaI-digested pBlueScript II SK (+) vector. The final construct was microinjected into *Drosophila* embryos (BL78781 stock) to generate the knockout strain.

### *Drosophila* sleep analysis

Adult *Drosophila* cultured at 29℃ for 3 or 10 days were packed into 65 mm × 5 mm glass motion tubes containing 5% sucrose AGAR food. The experiments were carried out in an incubator with a temperature of 25±1℃ and a relative humidity of 50%±5%. Turn on the lights at ZT0 (09:00 local time) and turn off the lights at ZT12 (21:00 local time). The *Drosophila* were always given at least 1 day of acclimatization before sleep analysis. Sleep and motor activity data were collected in 1-minute boxes using the *Drosophila* Activity Monitoring System (TriKinetics). Sleep was defined as periods of 5 min or more of behavioral immobility. Sleep/activity parameters (total sleep, mean sleep episode duration, and number of sleep episodes) were analyzed for each 24 hours and averaged across 2 or more days (excluding the first day to allow for acclimation). Sleep analysis was conducted using the MATLAB program SCAMP, as described previously.

### Overexpression of circRNA-mediated BT proteins in *Drosophila*

The *Drosophila* strain *elav*-GAL4 was purchased from the Tsinghua Fly Center. The backward ORF region of *dm*.circROLS, *dm*.circSK, *dm*.circKUZ, *dm*.circFZ, and *dm*.circTRPM were synthesized in the forward orientation, then cloned into the pUAST-attBVector plasmid at the multiple cloning site (**Fig. 6C**). Following plasmid extraction, the construct was injected into *Drosophila* embryos (pole cells). After eclosion, the *Drosophila* were crossed with background or balancer chromosomes *Drosophila* to achieve homozygosity, resulting in the UAS-*dm*.circROLS (Ⅱ), UAS-*dm*.circSK(Ⅱ), UAS-*dm*.circKUZ (Ⅲ), UAS-*dm*.circFZ (Ⅱ) and UAS-*dm*.circTRPM (Ⅲ) homozygous *Drosophila* strain.

---

**Fig. S1.**
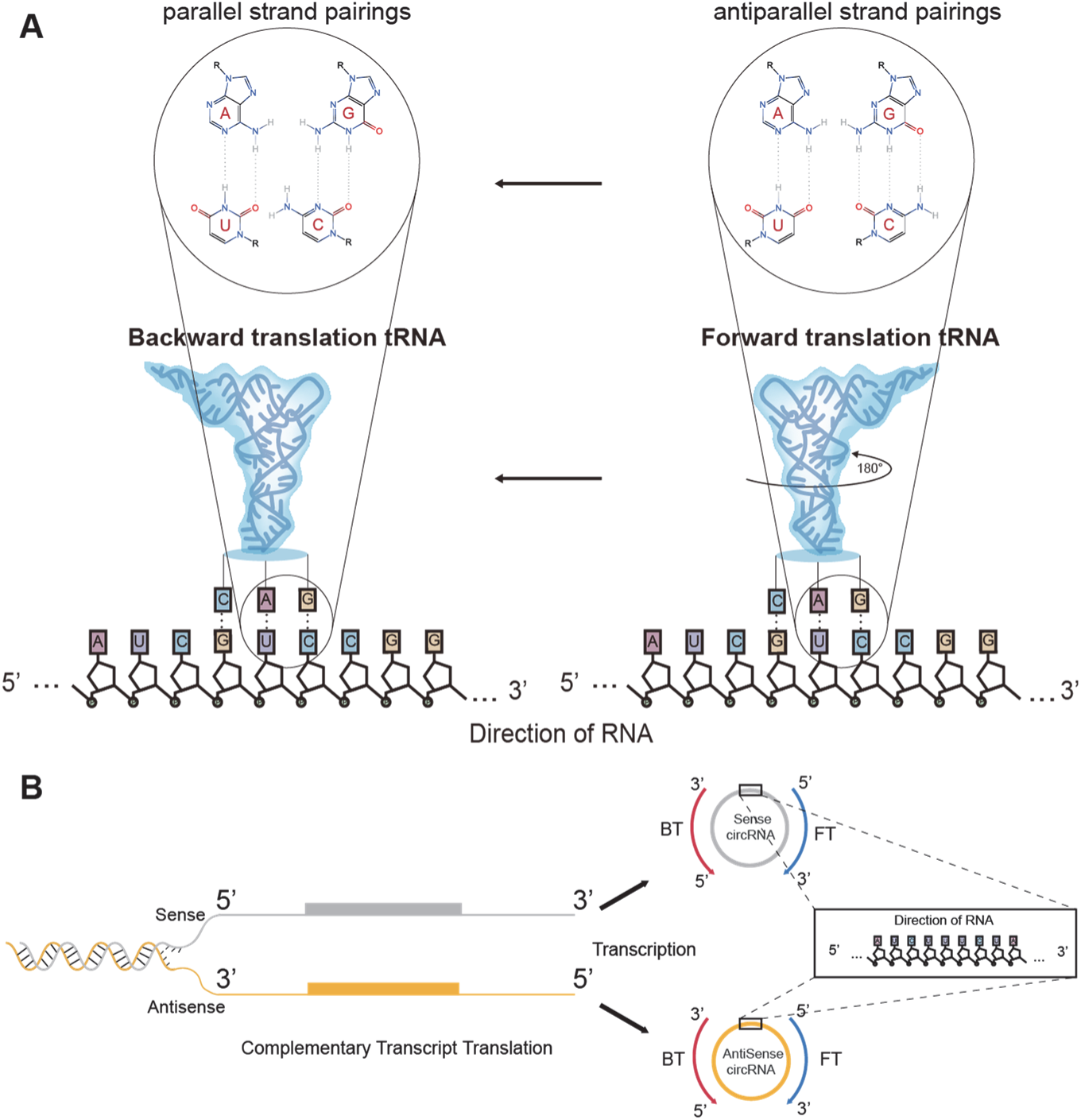
Schematic illustration of the relationship between backward/forward translation and complementary translation. **(A)** Conceptual diagram of reverse pairing of tRNAs to circRNA bases in a flipped orientation during backward translation. **(B)** Comparison between backward translation and translation of ‘antisense transcript’. Backward translation and forward translation can occur in circRNAs transcribed from both sense and antisense DNA strands.

**Fig. S2.**
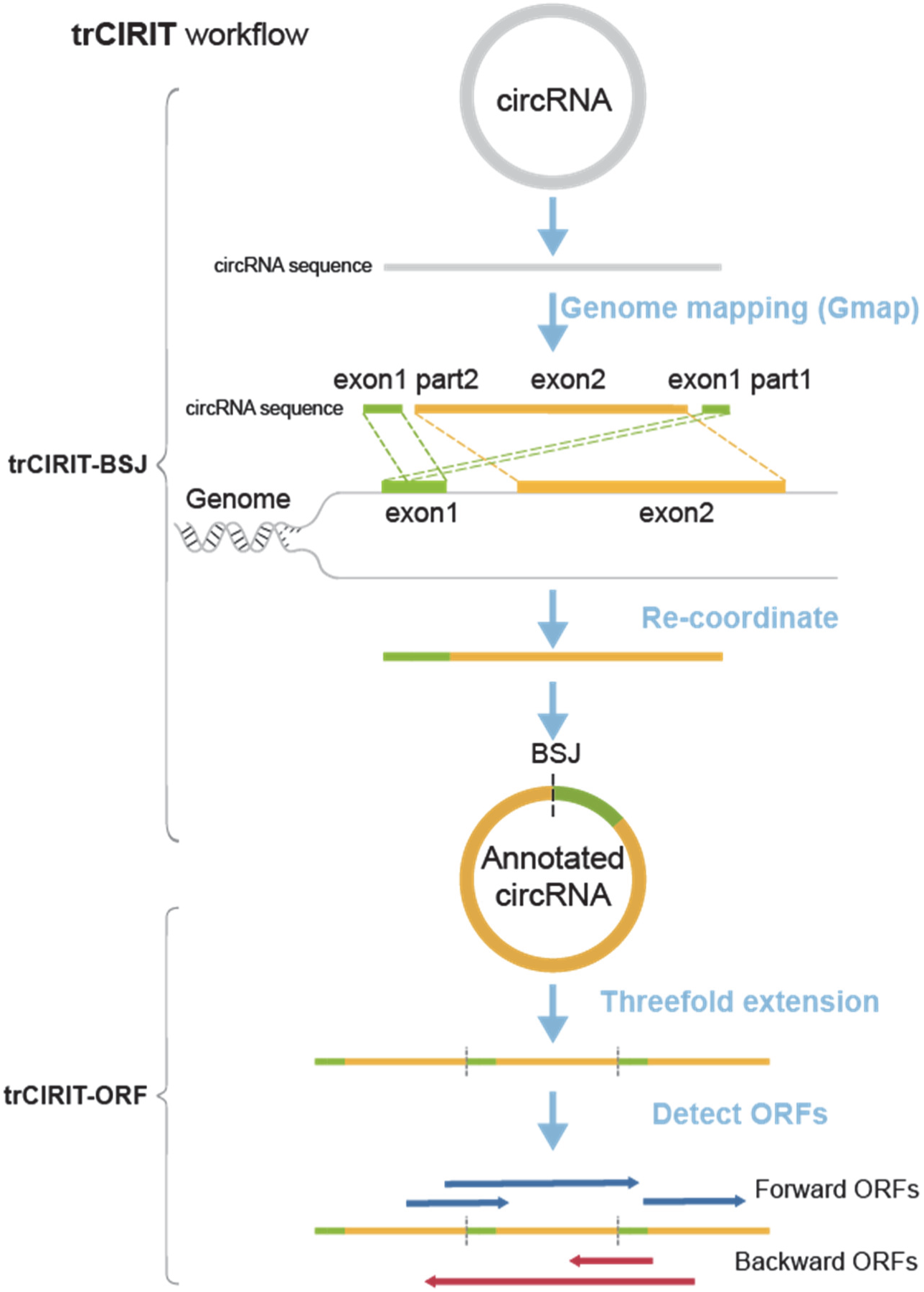
The trCIRIT workflow for circRNA annotation and ORF identification. The workflow consists of two main modules: trCIRIT-BSJ, which maps circRNA sequences to the genome, re-coordinates exon positions, and annotates back-splice junctions (BSJ); and trCIRIT-ORF, which extends circRNA sequences threefold and detects both forward and backward ORFs to characterize potential translational features of circRNAs.

**Fig. S3.**
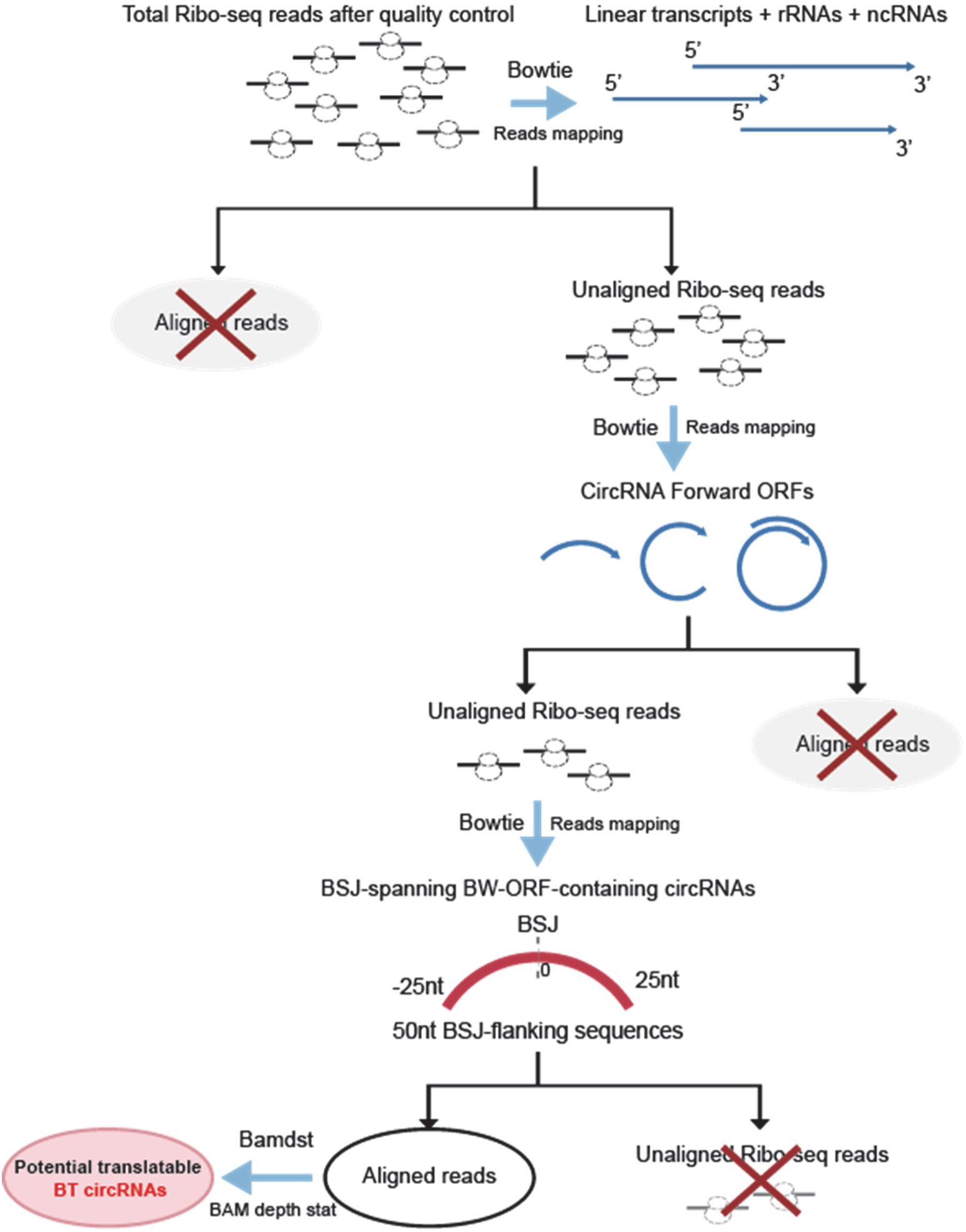
Bioinformatics workflow to detect active backward translation using Ribo-seg evidence. After quality control, reads aligned to rRNAs, linear mRNAs (GRCh38.p14), ncRNAs (RNAcentral), and circRNA forward ORFs were sequentially removed. The remaining reads were aligned exclusively to the back-splice junction (BSJ) and its flanking 25 nt sequences (a total of 50 nt) spanned by backward ORFs using Bowtie v1.0.0 (-v 1). BSJ-spanning reads were identified as evidence of potential backward translation activity.

**Fig. S4.**
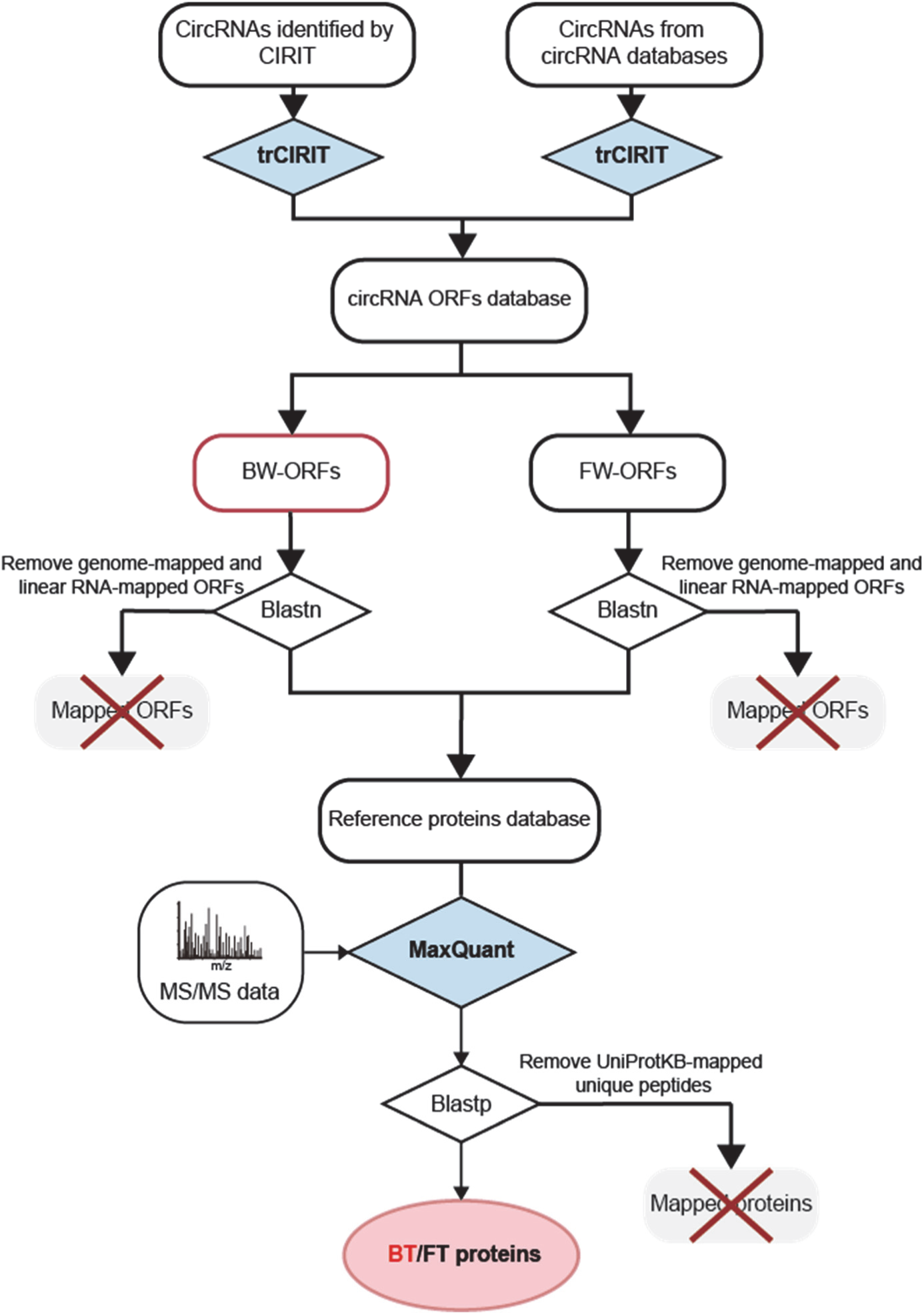
Bioinformatic workflow to detect backward translation products using mass spectrometry data. CircRNAs identified by trCIRIT or external databases are used to construct a circRNA ORF database. Forward (FW) and backward (BW) ORFs are filtered to remove those mapping to the genome or linear RNAs. MS/MS spectra are analyzed with MaxQuant, and unique peptides not matching UniProtKB are retained as evidence of potential backward translation (BT) or forward translation (FT) circRNA-derived proteins.

**Fig. S5.**
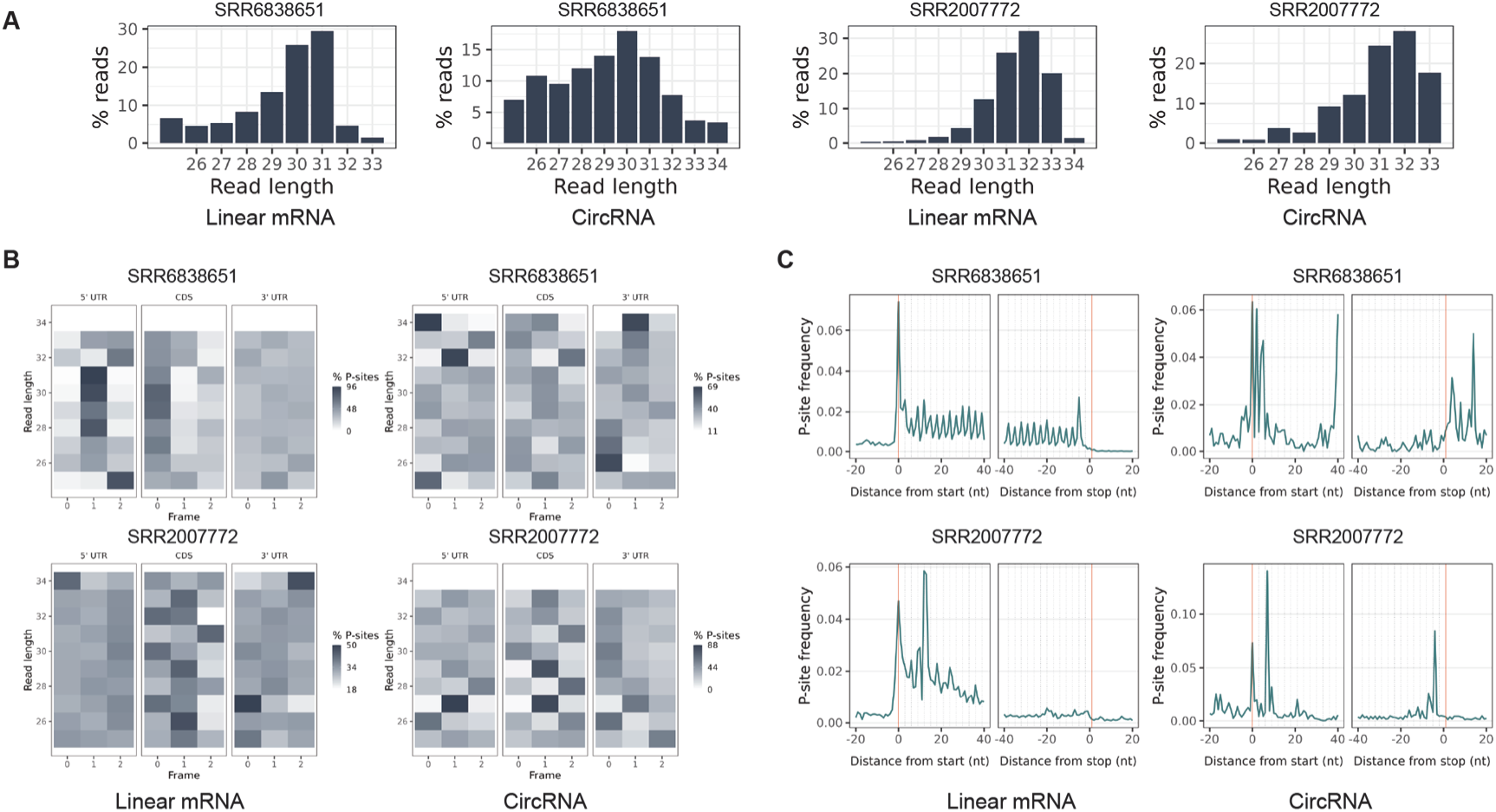
Diagnostic analysis of ribosome profiling for backward translation. **(A)** Read length distribution of linear transcripts and backward-translated circRNAs. **(B)** Periodicity of reads from linear transcripts and backward-translated circRNAs. **(C)** Ribosome P-site distribution frequency of linear transcripts and backward-translated circRNAs. For circRNA analysis, to facilitate plotting, we reversed both the backward ORFs and the reads used for alignment. The backward ORFs were converted to the normal 5’-3’ reading sequence (i.e., from GAT/AAT/AGT…GTA to ATG…TGA/TAA/TAG).

**Fig. S6.**
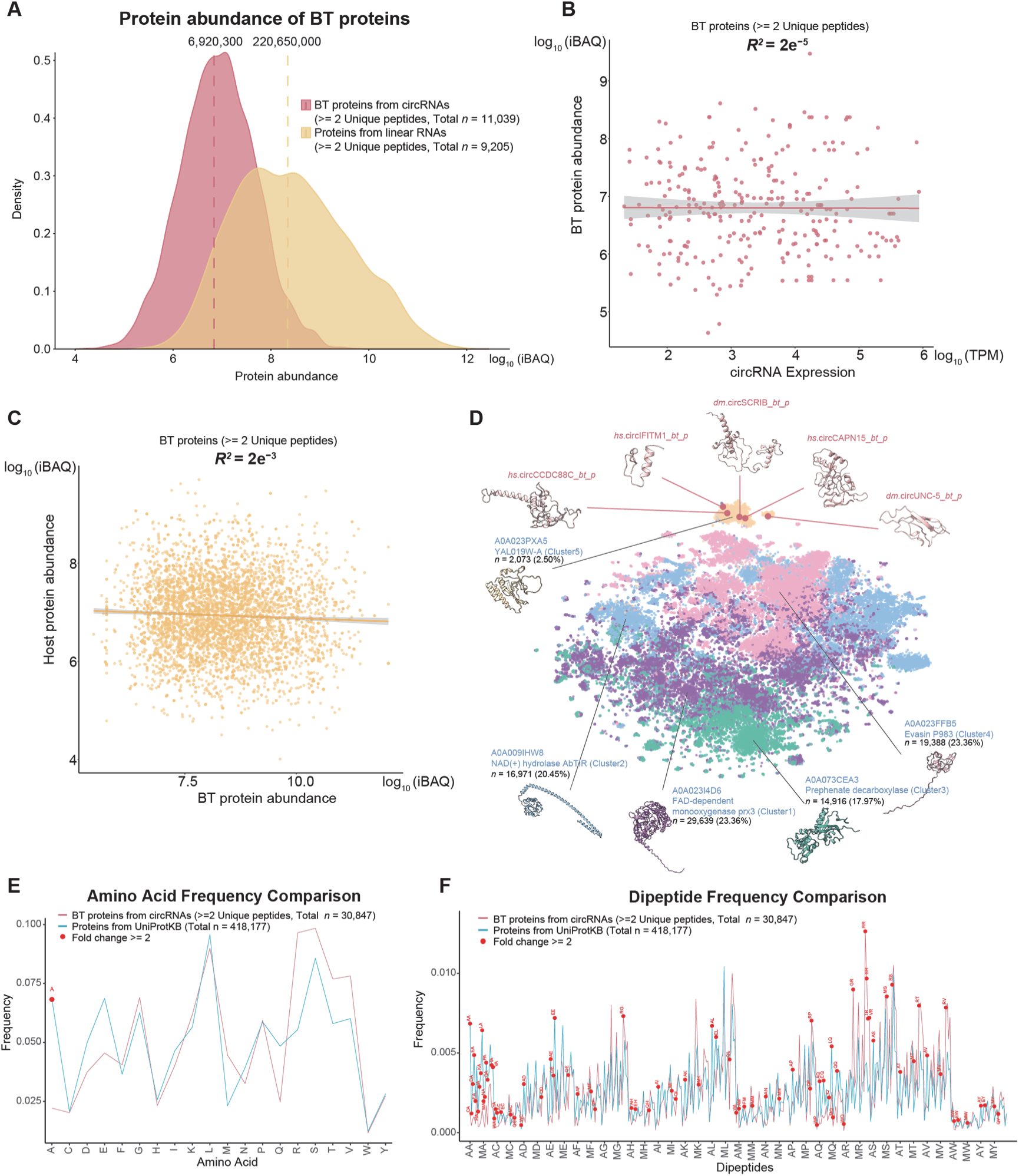
Expression characteristics of backward translation (BT). **(A)** Abundance comparison between BT proteins and the proteins coded by linear mRNAs (sourced from the UniProtKB). **(B)** Abundance correlation between circRNAs and the BT proteins they produced. The correlation analysis was demonstrated on a parallel transcriptome/proteome dataset determined on 70 liver samples (dataset ID: PXD006512). Sample size: 279 BT proteins corresponded to 239 parental circRNAs. **(C)** Abundance correlation between BT proteins and their counterpart host proteins. The linear regression analysis was utilized, and the coefficient of determination (*R*^2^) was calculated. Sample size: 5,413 BT proteins corresponded to 2,187 parental proteins. **(D)** Structural features of BT proteins. The structure of five selected BT proteins, including three human BT proteins (*hs*.circCCDC88C_*bt*_*p*, *hs*.circIFITM1_*bt*_*p*, and *hs*.circCAPN15_*bt*_*p*) and two *Drosophila* BT proteins (*dm*.circSCRIB_*bt*_*p* and *dm*.circUNC-5_*bt*_*p*), were modeled with AlphaFold and subsequently optimized with 50ns molecular dynamics with Gromacs, adopting the GROMOS force field and the TIP3 water environment. Structural comparison between five selected BT proteins and the present known proteins, including 542,337 AlphaFold-modeled structures. For large-scale comparison, the structures were first represented as the average embedding tokens with the ProtT5-XL-U50 model of ProtTrans. The K-means method was then utilized to cluster the structures. The t-SNE method of the cuML package, Matplotlib, and Seaborn were utilized to visualize the clustering result. **(E)** Comparison of amino acid usage frequencies between standard proteins in UniProt and BT protein. **(F)** Comparison of dipeptide frequencies between BT protein and standard proteins in UniProt.

**Fig. S7.**
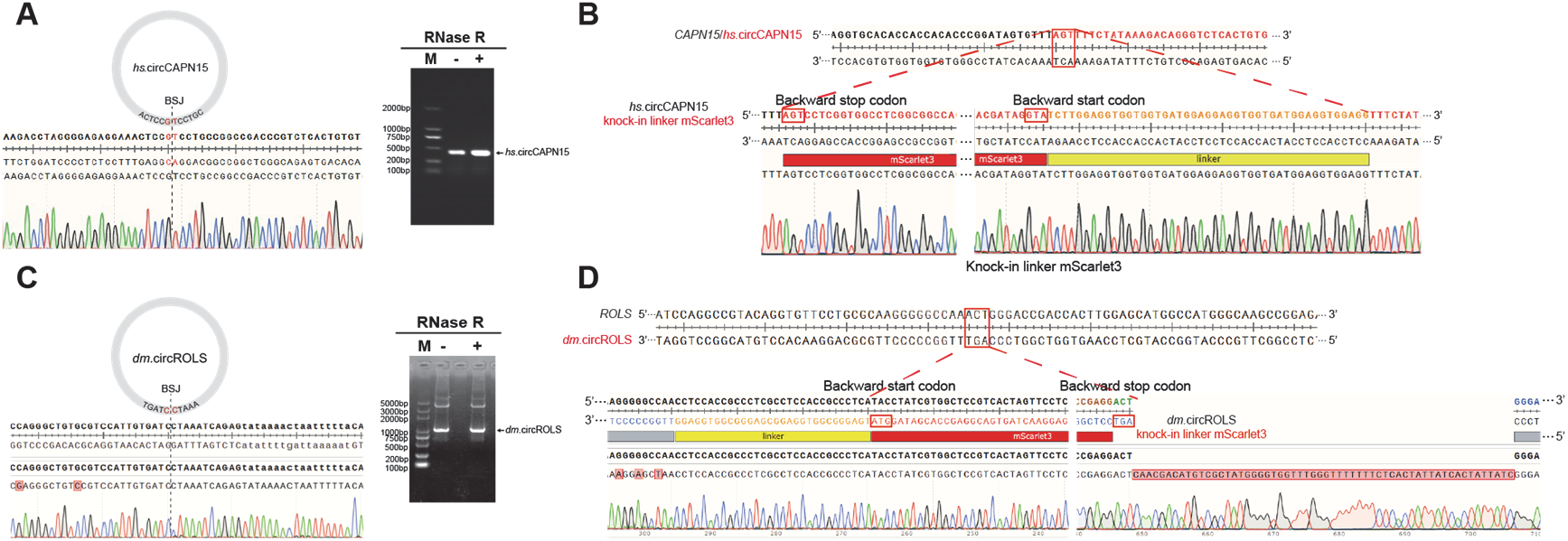
Validation of endogenous circular RNA expression and knock-in (KI) construct. **(A)** RT-PCR validation of *hs.*circCAPN15. The presence of a 329 bp band following RNase R treatment corrects circularization and expression of the circular RNA *hs.*circCAPN15. **(B)** Sanger sequencing verification of the *hs.*circCAPN15 knock-in in HeLa cells. **(C)** RT-PCR validation of *dm.*circROLS. The 963 bp band retained after RNase R treatment, along with Sanger sequencing results, correct circularization and expression of *dm.*circROLS. A schematic of the plasmid used for the knock-in experiment is shown. **(D)** A reverse ORF encoding the red fluorescent protein mScarlet3 was inserted immediately upstream of the stop codon (5’-AGT-3’) in the backward ORF of *dm.*circROLS. The knock-in was performed using CRISPR/Cas9 genome editing.

**Fig. S8.**
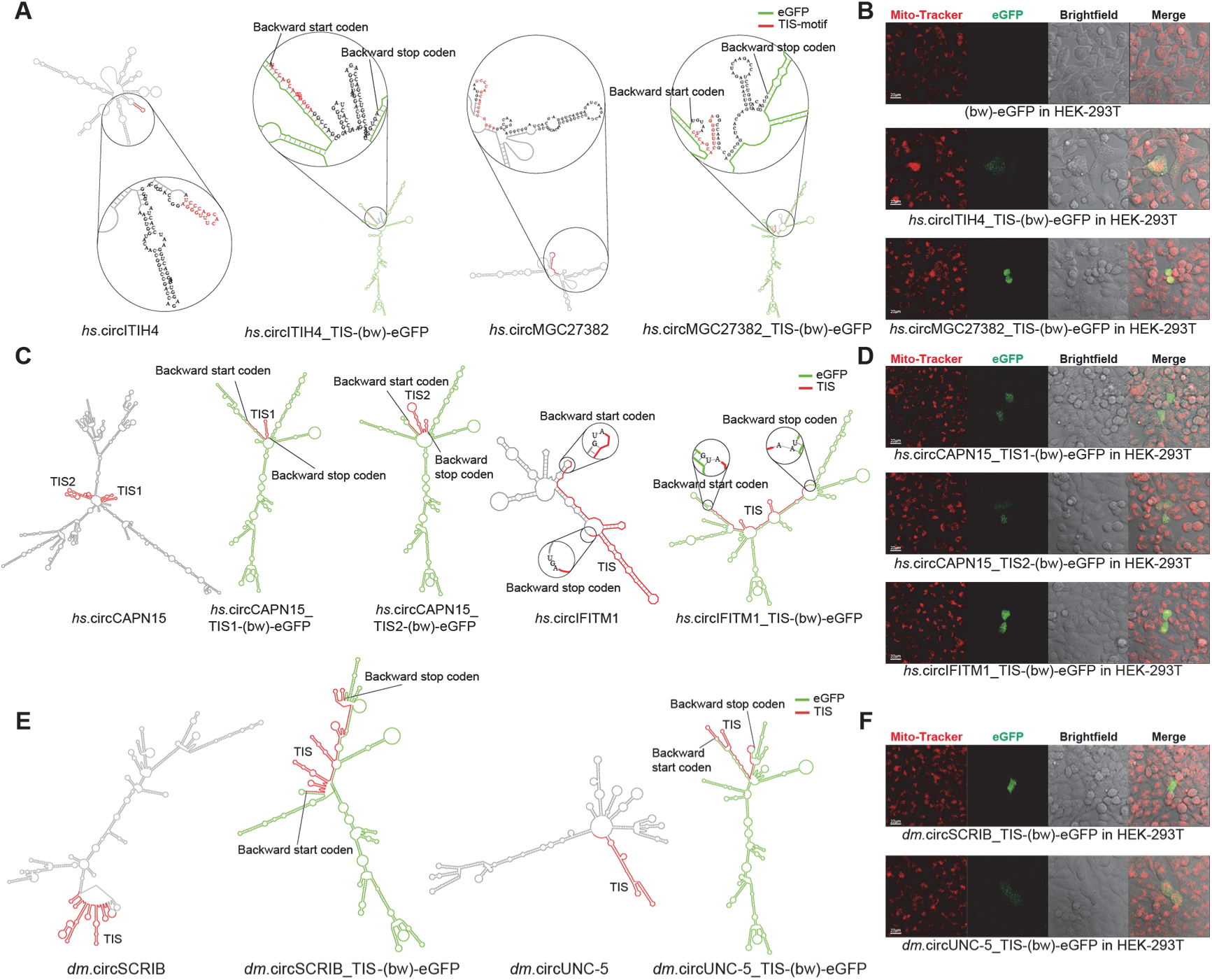
Validation of translation initiation site (TIS) sequences for initiating backward translation. **(A)** Extraction of candidate translation initiation site (TIS) sequences from *hs.*circITIH4 and *hs.*circMGC27382. Each TIS was inserted into engineered circRNAs, linking the start and stop codons of the reverse eGFP open reading frame (ORF). Both TISs contained the conserved ‘TCCCA…TGGGA’ motif but failed to form stem-loop structures when inserted into the circ-(bw)-eGFP constructs. **(B)** Representative confocal microscopy images showing eGFP fluorescence in HEK-293T cells transfected with circ-(bw)-eGFP constructs containing no TIS, *hs.*circITIH4-TIS-(bw)-eGFP, or *hs.*circMGC27382-TIS-(bw)-eGFP. The eGFP fluorescence is shown in green. Scale bar, 20 μm. A fluorescent signal was detected in approximately 1-2 out of every 100,000 cells. **(C)** Extraction of TISs from *hs.*circCAPN15 (two candidates) and *hs.*circIFITM1. These TISs did not contain the canonical “TCCCA…TGGGA” motif but were capable of forming stem-loop structures upstream of the eGFP ORF. Each was inserted into circRNAs to generate circ-(bw)-eGFP constructs. **(D)** Confocal images showing eGFP fluorescence in HEK-293T cells transfected with *hs.*circCAPN15-TIS1-(bw)-eGFP, *hs.*circCAPN15-TIS2-(bw)-eGFP, or *hs*.circIFITM1-TIS-(bw)-eGFP. eGFP signals are shown in green. Scale bar, 20 μm. A fluorescent signal was observed in ∼1-2 out of 100,000 cells. **(E)** Extraction of TISs from *dm.*circSCRIB and *dm.*circUNC-5. **(F)** Confocal microscopy of HEK-293T cells transfected with *dm.*circSCRIB-TIS-(bw)-eGFP or *dm.*circUNC-5-TIS-(bw)-eGFP constructs. All engineered circRNAs led to detectable eGFP fluorescence. Representative images are shown; eGFP in green. Scale bar, 20 μm. Fluorescent signal was again observed in approximately 1-2 out of every 100,000 cells. After 24 h, live cells were stained with Mito-Tracker (red, mitochondria).

**Fig. S9.**
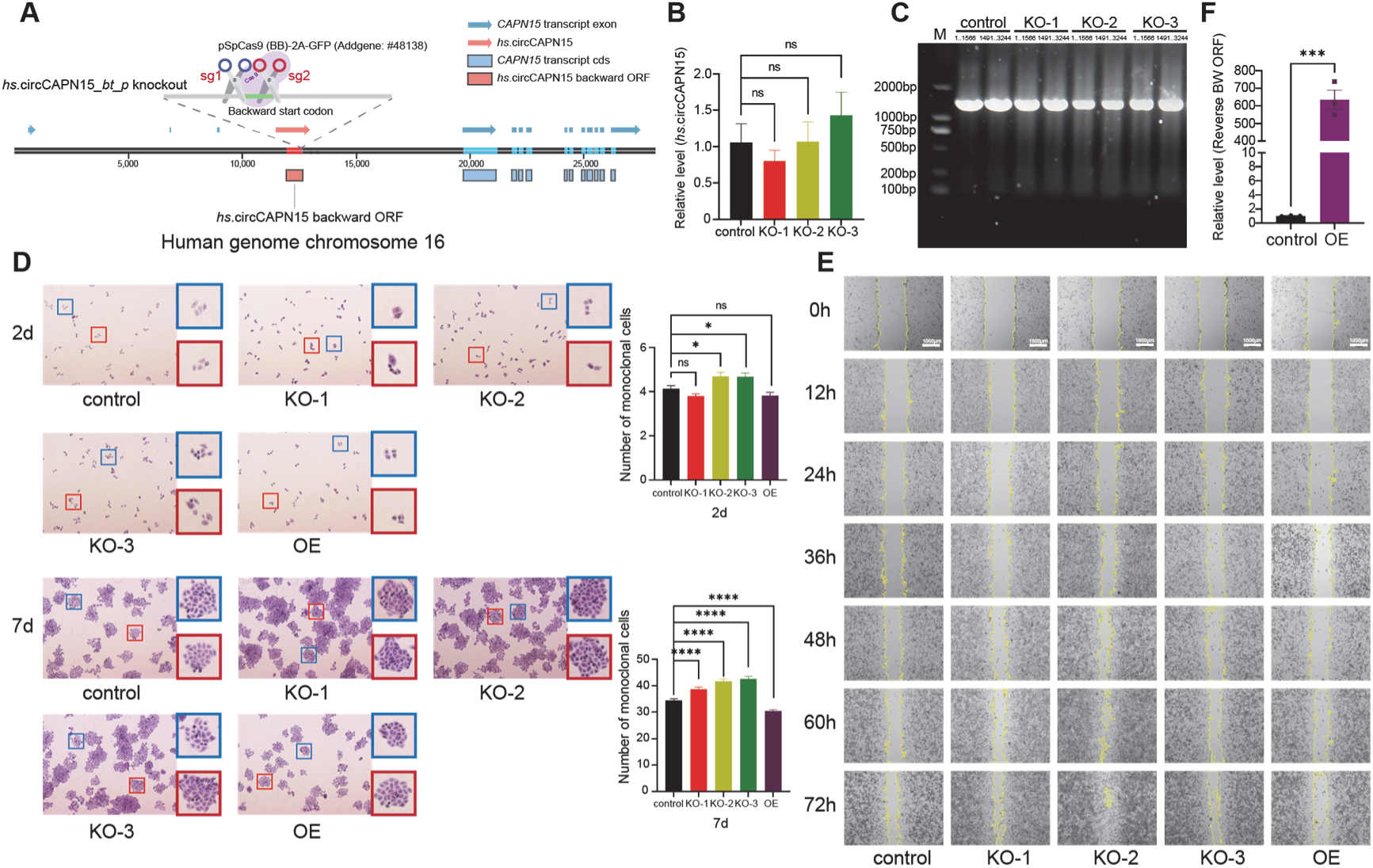
Functional characterization of the BT protein *hs.*circCAPN15*_bt_p* in human cells. **(A)** Schematic representation of the genomic organization of *hs.*circCAPN15 and its host gene *CAPN15*. The *hs.*circCAPN15 circRNA is transcribed from a non-coding intronic region of *CAPN15*, spatially separated from the protein-coding sequence (CDS). **(B)** RT-qPCR analysis of *hs*.circCAPN15 expression before and after CRISPR/Cas9-mediated knockout (KO). Statistical comparisons between groups were performed using Mann-Whitney U (GraphPad Prism 9.2.0) with significance levels indicated as follows: “*”: *p* <0.05; “**”: *p* <0.01; “***”: *p* <0.001; “****”: *p* <0.0001; *ns*: not significant (*p*>0.05) (*n*=3). **(C)** RT-PCR analysis of the linear *CAPN15* transcript before and after *hs.*circCAPN15 knockout. The coding sequence (CDS) of the dominant transcript variant *CAPN15*-201 (3,261 bp) was amplified in two overlapping fragments (nucleotides 1–1566 and 1491–3244). **(D)** Plate colony formation assay assessing the impact of *hs.*circCAPN15*_bt_p* KO and overexpression (OE) on HeLa cell proliferation at 2 and 7 days post-treatment. Representative fields were marked with red and blue for visual cell counting. **(E)** Cell migration assay (scratch wound healing) measuring the effects of *hs.*circCAPN15*_bt_p* KO and OE on HeLa cell motility. Scratch closure was recorded every 12 hours up to 72 hours. **(F)** RT-qPCR quantification of the forward-oriented linear *hs.*circCAPN15*_bt_p* ORF expression before and after overexpression treatment. Bar graphs show mean ±SEM for biological replicates (*n* =3). Statistical significance: *ns*: not significant, “*”: *p* <0.05; “**”: *p* <0.01; “***”: *p* <0.001; “****”: *p* <0.0001.

**Fig. S10.**
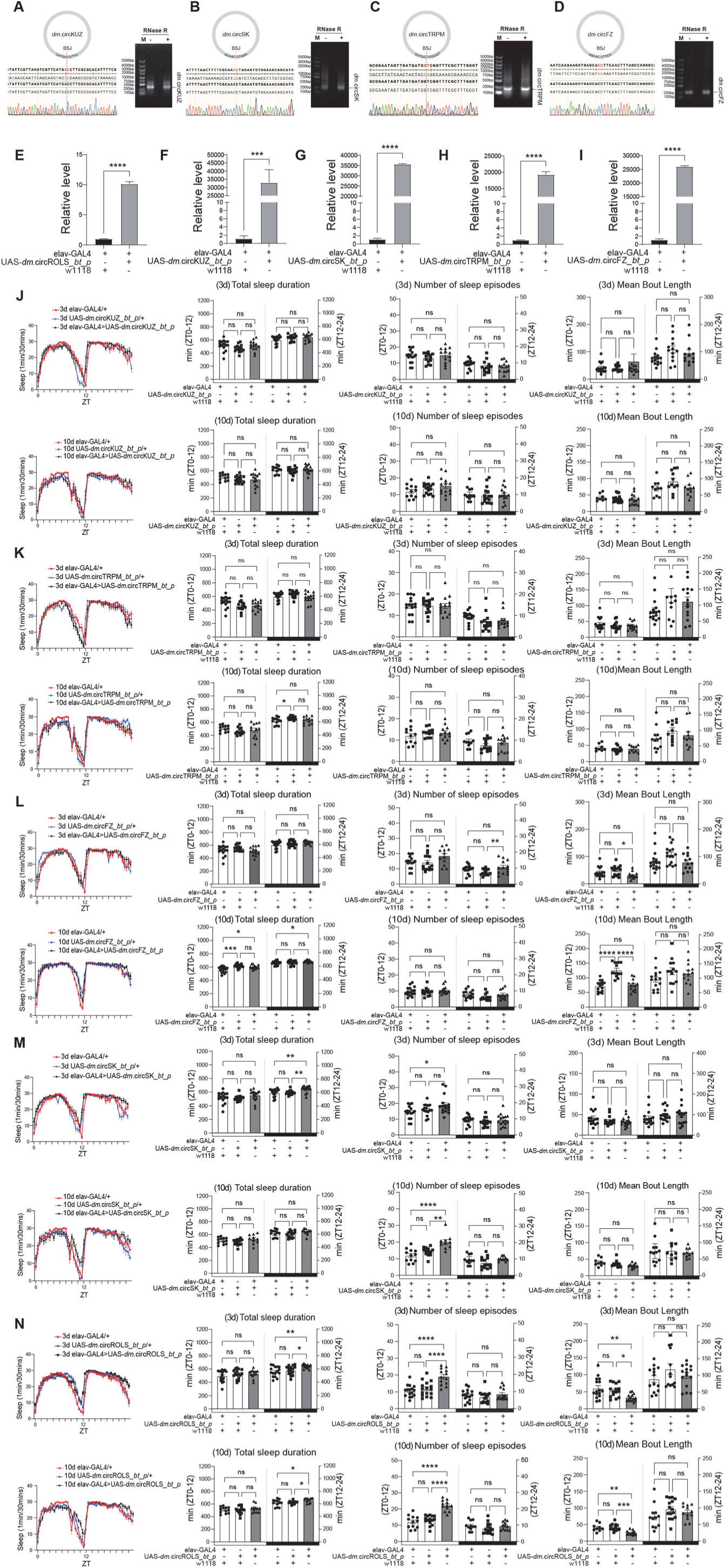
Screening of sleep-related BT circRNAs in *Drosophila*. **A-D,** Predicted secondary structures and RT-PCR validation of candidate brain-translated circRNAs: *dm.*circKUZ (A), *dm.*circSK (B), *dm.*circTRPM (C), and *dm*.circFZ (D). For each, a distinct RT-PCR band was retained after RNase R treatment (354 nt for *dm.*circKUZ, 345 nt for *dm.*circSK, 208 nt for *dm.*circTRPM, and 232 nt for *dm.*circFZ), and Sanger sequencing confirmed the presence of the back-splice junction, validating circularity and expression. **E-I**, RT-qPCR quantification of overexpression levels for linearized BT-circRNA constructs: *dm.*circROLS*_bt_p* (E), *dm.*circKUZ*_bt_p* (F), *dm.*circSK*_bt_p* (G), *dm.*circTRPM*_bt_p* (H), and *dm.*circFZ*_bt_p* (I). **J-N**, Sleep phenotyping of 3-day-old and 10-day-old *Drosophila* with pan-neuronal overexpression of BT circRNAs: *dm.*circKUZ*_bt_p* (J), *dm.*circTRPM*_bt_p* (K), *dm.*circFZ*_bt_p* (L), *dm.*circSK*_bt_p* (M), and *dm.*circROLS*_bt_p* (N). Sleep profiles depict 30-minute binned sleep accumulation over 24-hour periods. Total sleep duration represents the summed sleep time. The number of sleep episodes indicates a sleep bout. Mean Bout Length represents the mean sleep episode duration. An increase in the number of sleep episodes accompanied by a decrease in mean bout length indicates a sleep fragmentation phenotype. Mann-Whitney U (GraphPad Prism 9.2.0) was used to statistically analyze the inter group differences in mean bout length, with the following significance levels: *ns*: not significant, “*”: *p* <0.05, “**”: *p* <0.01, “***”: *p* <0.001, “****”: *p* <0.0001; (*n*>10)White and black bars on the x-axis denote day (ZT0–12) and night (ZT12–24), respectively. Mann-Whitney U (GraphPad Prism 9.2.0) was used to statistically analyze the inter group differences in mean bout length, with the following significance levels: *ns*: not significant, “*”: *p* <0.05, “**”: *p* <0.01, “***”: *p* <0.001, “****”: *p* <0.0001; (*n*>10). The other statistical comparisons between groups were performed using unpaired two-tailed t-tests (GraphPad Prism 9.2.0). Statistical significance: *ns*: not significant, “*”: *p* <0.05, “**”: *p* <0.01, “***”: *p* <0.001, “****”: *p* <0.0001; (*n*>10).

**Fig. S11.**
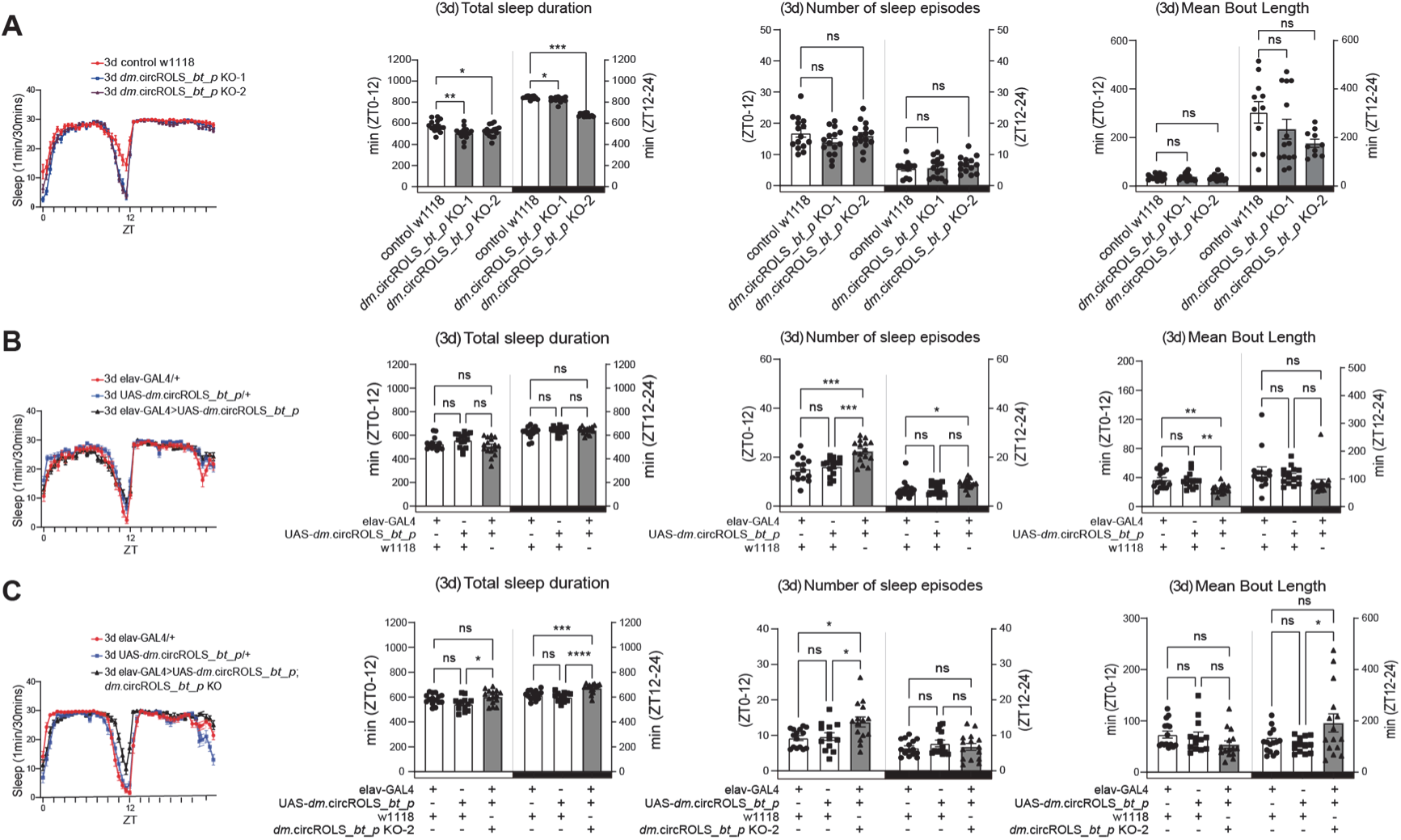
Overexpression of *dm*.circROLS*_bt_p* rescues sleep phenotype in KO *Drosophila*. **(A)** Sleep monitoring of a 3-day-old *dm.*circROLS*_bt_p* KO-1 and KO-2 *Drosophila* and w1118 (wild-type) control with pan-neuronal driver. **(B)** Sleep monitoring of 3-day-old *Drosophila* with pan-neuronal overexpression of *dm.*circROLS*_bt_p* and control groups (*elav*-GAL4/+, UAS-*dm.*circROLS*_bt_p*). **(C)** Sleep monitoring of 3-day-old *Drosophila* with pan-neuronal overexpression of *dm.*circROLS*_bt_p* in the KO-2 background and control (*elav*-GAL4/+, UAS-*dm.*circROLS*_bt_p*). **A-C**, Sleep profiles depict 30-minute binned sleep accumulation over 24-hour periods. Total sleep duration represents the summed sleep time. The number of sleep episodes indicates a sleep bout. Mean Bout Length represents the mean sleep episode duration. An increase in the number of sleep episodes accompanied by a decrease in mean bout length indicates a sleep fragmentation phenotype. White and black bars along the x-axis indicate day (ZT0–12) and night (ZT12–24), respectively. Mann-Whitney U (GraphPad Prism 9.2.0) was used to statistically analyze the inter group differences in mean bout length, with the following significance levels: *ns*: not significant, “*”: *p* <0.05, “**”: *p* <0.01, “***”: *p* <0.001, “****”: *p* <0.0001; (*n*>10). The other statistical comparisons between groups were performed using unpaired two-tailed t-tests (GraphPad Prism 9.2.0). Statistical significance: *ns*: not significant, “*”: *p* <0.05, “**”: *p* <0.01, “***”: *p* <0.001, “****”: *p* <0.0001; (*n*>10).

